# Highly specific mRNA cleavage by the MazF endoribonuclease orchestrates stationary transcriptome remodeling and rapid regrowth in Gram-positive bacteria

**DOI:** 10.64898/2026.08.06.743204

**Authors:** Regev Frenkel, Shira Omer, Tom Borenstein, Bar Tenenbaum, Shani Shalev, Nadejda Sigal, Yuyi Li, Polina Guler, Avigail Zarzar, José R Penadés, Avigdor Eldar

## Abstract

Bacterial toxin-antitoxin (TA) systems are classically viewed as stress-activated toxic switches. Specifically, ribonucleolytic toxins are thought to indiscriminately cleave RNA to halt cellular growth. We recently showed that the MazF toxin of *Bacillus subtilis* targets an unusually strict 6bp RNA cleavage sequence, but the implications of this stringent specificity were unknown. Here, we demonstrate that the MazEF system functions as a non-lethal post-transcriptional regulator in *B. subtilis*. Using a specialized single cell fluorescent reporter and transcriptome profiling, we show that MazF is uniformly activated across the population upon entry into the stationary phase, where it cleaves a narrow mRNA regulon to reshape gene expression. Rather than inhibiting growth, MazF activation tunes down the Spo0A stress response by repressing the mRNA level of its kinases. Reduced stress leads to an adaptive shortening of the lag phase upon nutrient replenishment. Furthermore, MazEF’s structural architecture, cleavage specificity, and impact on growth recovery are highly conserved across Gram-positive bacteria. Altogether, our findings redefine a paradigmatic toxin as a precision global mRNA stress regulator that primes cells for rapid regrowth.

**Graphical abstract:** 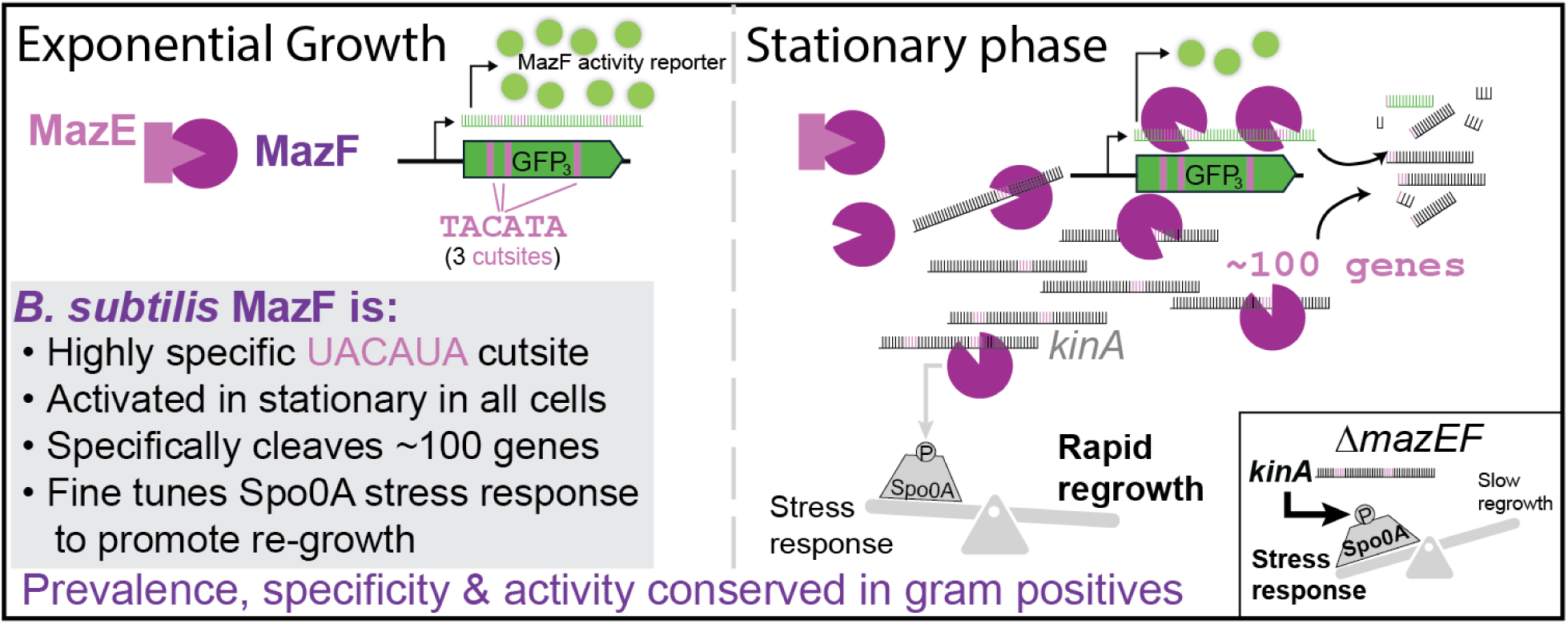

## 1. Introduction

Toxin-Antitoxin (TA) systems are highly prevalent in bacteria, typically comprising a protein toxin that can halt cellular growth and an antitoxin that blocks this activity(1–5). TA systems are primarily characterized as tightly regulated switches that induce dormancy or death under severe stress. Historically, they were shown to serve a role in mobile elements maintenance via post-segregation killing(2, 6–10) and maybe also as modulators of persistence(2, 11–14). More recently, their prominence as anti-phage defense mechanisms have been well demonstrated(15, 16).

An alternative, yet highly debated, hypothesis posits that certain TA systems function not as lethal switches, but as fine-tuned post-transcriptional regulators of cellular physiology(17, 18). However, validating this regulatory role has been a major challenge in the field. Many purported physiological effects remain contested because studies frequently rely on artificial toxin overexpression, lack genetic complementation, or fail to demonstrate the endogenous *in vivo* activity of the toxin under physiological conditions (5, 19, 20).

The MazEF system, a paradigmatic Type II TA system where the MazF toxin acts as an endoribonuclease is prevalent in many bacterial species and its role is still highly contraversial(21, 22). *E. coli* MazF recognizes a core 3-base pair (3bp) RNA motif (5’-^ACA-3’) with additional fine tuning of neighboring bases(23–25). This low specificity leads to multiple cleavages of most cellular transcripts. Consequently, the *E. coli* MazEF system has been proposed to function in programmed cell death(26), persister formation(27), Ribosomal reorganization(28) and phage defense(29, 30). However, most of these effects have been strongly contested(2, 24, 25, 31–33). Interestingly, it was recently suggested that MazF resolves R-loop formation on the 16S-gene during transcriptional stress in a non-lethal manner(34). This benefit overcome the cost of low specificity cleavage of mRNAs.

In contrast with *E. coli*, we and others have recently shown that the main *B. subtilis* MazF homolog (coded by the gene *ndoA*) has a striking degree of sequence specificity, targeting a strict 6bp 5’-U^ACAUA-3’ motif(35, 36). This high level of specificity suggest that MazF serves a different function in the two species. Furthermore, while the MazEF system is not found in all *E. coli* strains, it is part of the *Bacillus* genus core genome(35). This widespread conservation of a highly precise, exact-match ribonuclease strongly implies a dedicated, fundamental role in mRNA regulation, in addition to the specialized interactions with arbitrium-controlled phages, we and others have recently described(35, 37–39).

To determine whether this highly specific endoribonuclease serves a non-toxic regulatory function, we developed genetic and single-cell tools to directly detect endogenous MazF activity *in vivo*. Using these approaches, we uncover a population-wide activation of MazF during the transition into the stationary phase. Rather than inducing lethality, MazF reshapes the transcriptome by cleaving a specific mRNA regulon, including transcripts linked to the Spo0A stress-response pathway. We demonstrate that this targeted RNA decay tunes down cellular Spo0A-dependent stress response, thereby shortening the lag phase and promoting a rapid return to growth upon nutrient replenishment.

Extending this analysis beyond *B. subtilis*, we find that MazEF’s structural architecture, precise RNA cleavage specificity, and physiological impacts on growth dynamics are broadly conserved among Gram-positive bacteria. Together, these findings redefine the role of a paradigmatic TA system, demonstrating that bacteria utilize sequence-specific mRNA cleavage as a global, non-lethal regulatory strategy to navigate environmental transitions.

## 2. Results

### A single-cell fluorescent reporter detects endogenous, sequence-specific MazF endoribonuclease activity

To probe the activity of MazF at different physiological conditions at the single cell level, we designed a fluorescent reporter for MazF activity. Our group and others have shown that the MazF RNA cleavage motif in *B. subtilis* is the 6bp sequence UACAUA. We constructed a set of 4 IPTG-inducible *sfGFP* reporters containing between 0 to 3 MazF cut-sites, designated GFP_i_, where i stands for the number of cleavage sites (0-3, Fig. 1a). We reasoned that if MazF is activated, the *sfGfp_0_* transcript would not be cleaved, while *sfGfp_1-3_* transcripts would be cleaved at a rate dependent on the number of cleavage sites and the activity of MazF, leading to reduction in GFP expression compared to the gfp_0_ reporter. As the modifications are silent, we expected these to not have a substantial effect in the absence of MazF activity.

**Figure 1:**
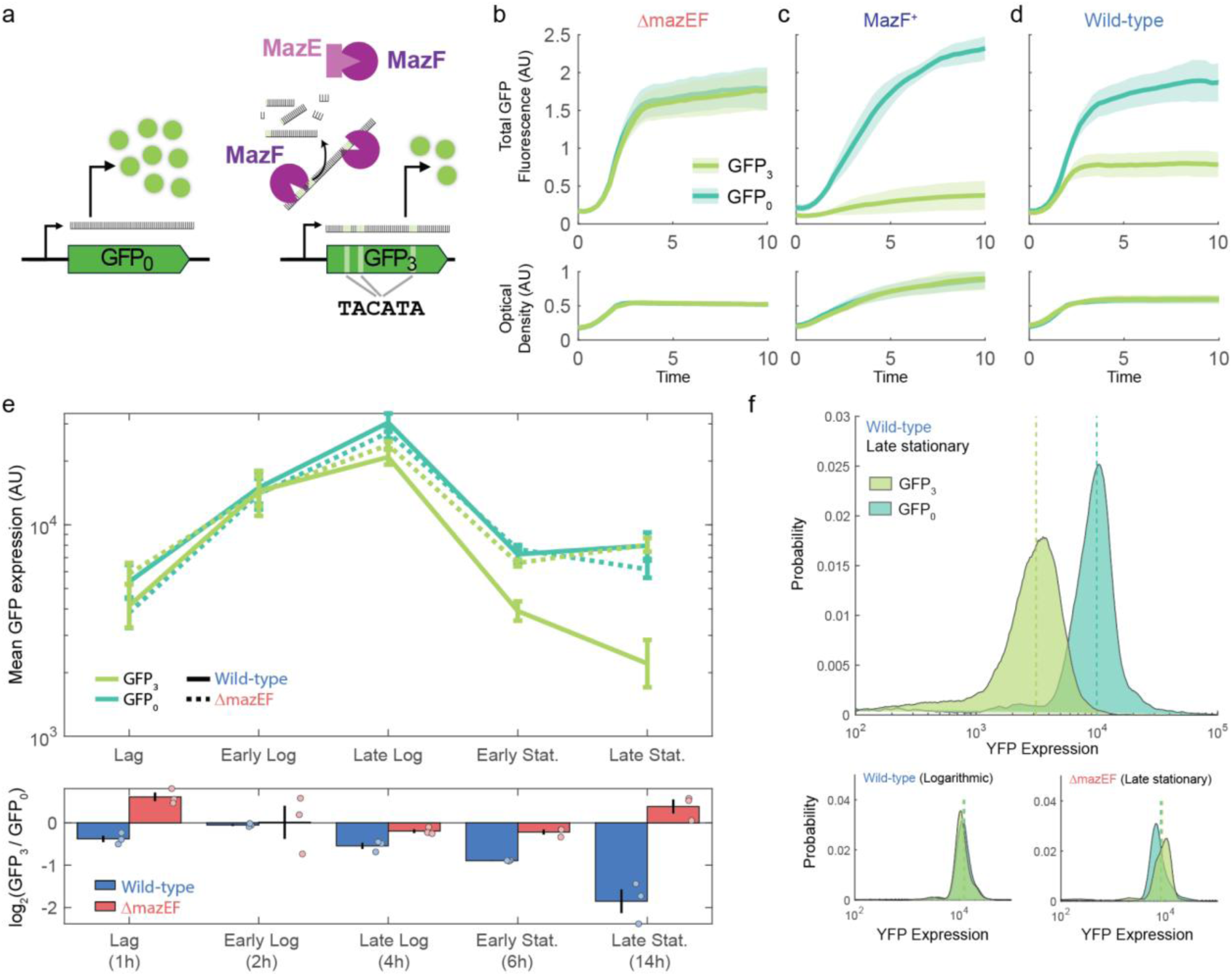
A fluorescent reporter for MazF activity reveals differential MazF activation along the population growth cycle. (a) Scheme of the GFP_i_ reporters. GFP_0_ has no MazF cut-sites, while GFP_3_ has three synonymous MazF cut-sites. If MazF (purple) is active and not blocked by MazE (light purple), then it will cleave the GFP_3_ transcript, resulting in lower GFP expression and fluorescence. (b-d) GFP_0_ (light green) and GFP_3_ (dark green) expression in (b) Δ*mazEF*, (c) MazF^+^ and (d) wild-type genetic backgrounds. (e) top: GFP_0_ (light green) and GFP_3_ (dark green) reporter expression for 1 hour induction of the two constructs, ending at the indicated time after dilution. Shown are results for wild-type (solid line) and Δ*mazEF* (dashed line). Bottom: ratio between GFP_3_ and GFP_0_ reporter levels at the different times and strain backgrounds. (f) Flow cytometer histograms of the GFP_0_ (light green) and GFP_3_ (dark green) reporters at late stationary (top) and logarithmic growth (bottom left) in the wild-type and of late stationary in the Δ*mazEF* strain (bottom right).

To verify the sensitivity of this reporter system, we used a *B. subtilis* Δ*mazEF* mutant strain and its derivative expressing *mazF* under a xylose inducible promoter. In a previous study, we showed that low expression of MazF (with 0.001% xylose) has mild effects on growth, while enabling detection of MazF activity(35). In accordance with its proposed utility, we find that the GFP_3_ reporter showed the same fluorescence as the GFP_0_ reporter in the Δ*mazEF* mutant background, but a greatly diminished level when MazF was expressed (Fig. 1b). The GFP_1,2_ reporters showed an intermediate response to MazF (Supplementary Fig. S1a,b). We therefore continued to characterize MazF activity using the fluorescence cleavage ratio, defined as the ratio between GFP_3_ and GFP_0_ reporters expression.

### MazF is uniformly activated across the population during the transition to stationary phase

Next, we introduced the GPF_i_ reporters into a wild-type background. We grew these strains in a plate reader on LB medium with continuous expression of the GFP reporters and found that their GFP expression differed during the growth cycle, with levels of GFP_1-3_ decreasing compared to GFP_0_ during the stationary state (Fig. 1d, Supplementary Fig. S1c,d). To better probe the temporal pattern of MazF activation, we utilized the inducibility of the GFP_i_ constructs. We diluted an overnight culture of uninduced GFP_0_ and GFP_3_ reporter strains into fresh LB medium and then added IPTG at different times after dilution before measuring GFP fluorescence of single cells using flow cytometry. Our results (Fig. 1e) demonstrate a clear reduction of the GFP levels in the GFP_3_ strain compared to the GFP_0_ strain during the lag phase (0 hours) and during early and late stationary phases (5h,14h) while no clear effect was observed during the logarithmic stage (1,3 hours). In a Δ*mazEF* mutant background GFP_3_ and GFP_0_ showed similar fluorescence levels. Examining the distribution of expression levels of single cells in GFP_0_ and GFP_3_ strains suggest that the reduction in GFP levels at stationary state occurs across the cell population rather than in a specific sub-population (Fig. 1f).

In *B. subtilis*, the *mazEF* operon is localized upstream of the *rsb* stressosome operon. To verify that the impact of this deletion is not due to a polar mutational effect on the expression of these genes, we constructed a *mazEF* complementation strain, by introducing a copy of the *mazEF* system with its native promoter into the chromosomal *amyE* locus in the Δ*mazEF* mutant. We found that this complementation largely rescued the wild-type effect, ruling out polar effects as an explanation of our observation (Supplementary Fig. S1e).

#### Transcriptomic profiling reveals a highly specific, cleavage-dependent MazF mRNA regulon

As MazF becomes active across the entire population, it is possible to probe its effect on the bacterial transcriptome using RNAseq. To this aim, we grew wild-type and corresponding Δ*mazEF* mutant cells overnight, diluted them into fresh LB and performed RNA sequencing assay on RNA extracted at the same time points used for measuring GFP fluorescence (Fig. 1e, methods, Supplementary File S1).

First, we used the RNAseq data to probe the level of MazF activity near putative MazF cut-sites. The expression cleavage ratio is defined as the average reduction of expression in the wild-type vs. Δ*mazEF* mutant(24). We average the cleavage ratio as a function of distance from putative cut-site (Fig. 2a). Averaging over >500 MazF TACATA cleavage sites from CDS with sufficiently high expression level (methods), we find a clear dip in average expression centered near the TACATA site and extending ∼250bp to each side (Fig. 2a). In agreement with the fluorescence cleavage ratio of GFP_3_ and GFP_0_, the depth of the dip was substantial during the lag time (1 h) and at early (6 h) and late (14 h) stationary phases, while no apparent dip was observed at early log (2 h) and a very small dip was found at late log (4 h) stages. Similar analysis around the presumed uncleavable motifs TACAT(C/G/T) did not show any dip at any time, further strengthening the previous indications for MazF cut-site specificity (see more below)(35, 36). Local cleavage activity results are therefore in good agreement with the results obtained by the reporter system. We note that the activation pattern of MazF does not strongly reflect its expression patterns, as MazEF expression is always in the upper 20% of expressed genes.

**Figure 2:**
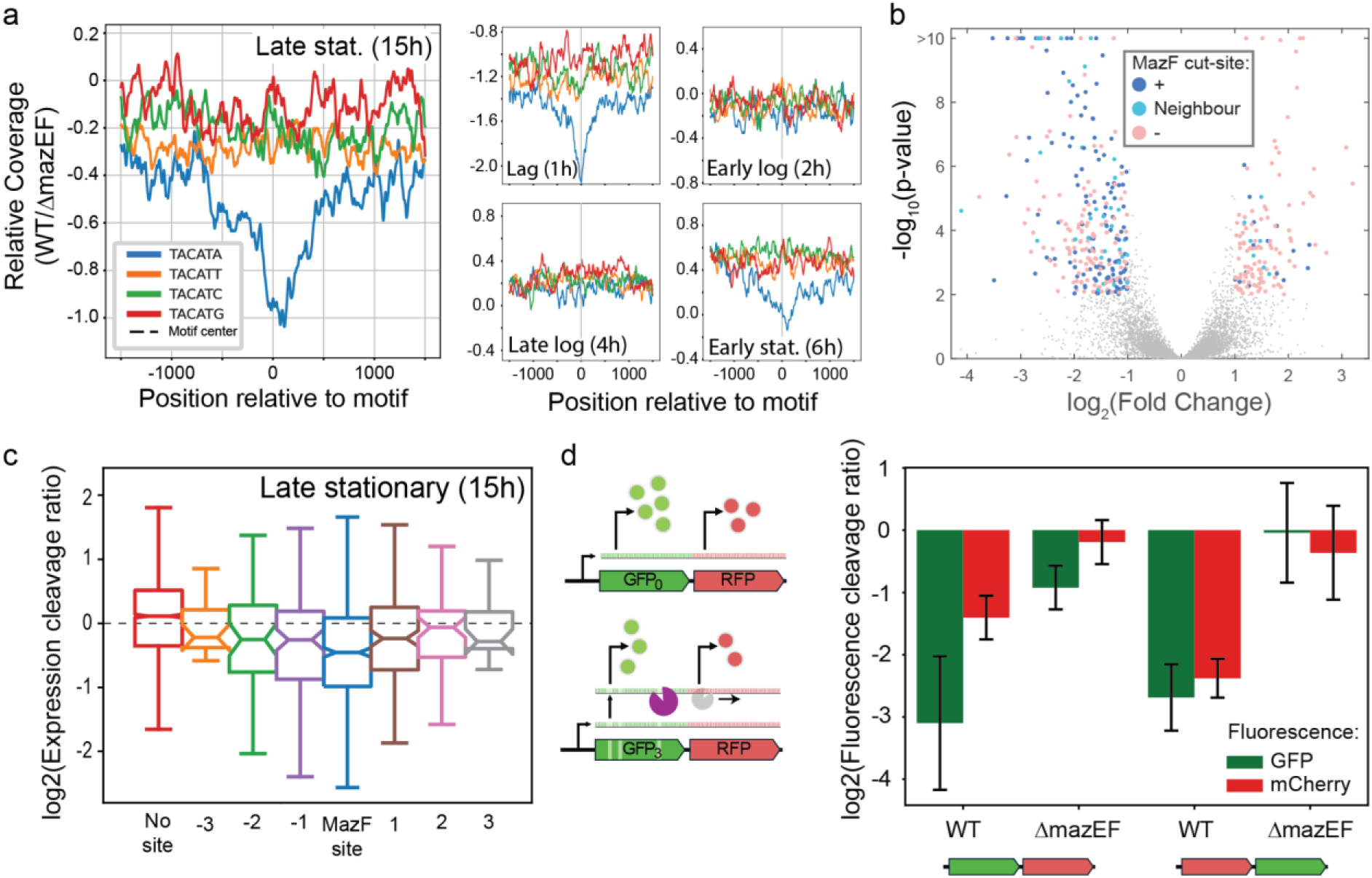
Comparative RNAseq between WT and Δ*mazEF* backgrounds reveals direct and indirect effects of MazF activity. (a) MazF cleavage ratio as a function of distance from a particular motif, averaged over all occurrences of this motif with sufficient expression level (methods). Results are shown for different growth stages (with cut-sites with sufficient expression level at different stages (methods). (b,c) Measures of expression cleavage ratio, defined as the fold change average gene expression ratio between wild-type and Δ*mazEF* strains. (b) Volcano plot of gene expression combined for the three conditions with significant MazF activity. Gray dots indicate all genes with expression cleavage ratio smaller than 2 or p-value>0.01. Colored markers highlight genes with a larger than 2 fold-change and p-value<0.01. Genes with MazF cut-site (dark blue), with a co-cistronic neighbour with a cut-site (light blue) and without a cut-site (red) are indicated. (c) Whisker plot of the logarithmic distribution of the expression cleavage ratio of different sets of genes; without a cut-site in their operon, or with varying gene distances from a co-cistronic gene with a cut-site according to legend. (d) Fluorescence cleavage ratio for *sfGfp* and a co-cistronic *mCherry* gene, defined as the relevant fluorescence ratio in a construct with GFP_0_ and GFP_3_. *mCherry* lacks MazF cut-sites. Two operon structures were measured; *mCherry-sfGfp* and *sfGfp-mCherry* in either WT or Δ*mazEF* backgrounds. All measurements were taken in the late stationary stage (14 hours past dilution) in LB medium.

Next, we wondered what is the effect of MazF on gene expression. We averaged the transcript expression over each gene in our background PY79 strain and studied the fold change in gene expression between the wild-type and the mutant at different times (Supplementary File S1). We found sets of 68 and 221 genes whose expression was reduced in the wild-type in any of the experiments by a factor of four or two, correspondingly. A smaller number of genes were found to be upregulated in the wild-type (22 and 128 for factors of 4 and 2) (Fig. 2b). Notably, genes coding for the MazF cut-site were strongly enriched in the set of genes with reduced fold change (40% in this set, compared with 15% of genes with absolute fold change smaller than 2). In accordance with this, we find that MazF cut-site coding genes showed on average a stronger reduction in gene expression in the wild-type compared to the Δ*mazEF* mutant than genes not coding for the MazF cut-site (Fig. 2c).

Interestingly, we also found that co-cistronic neighbours of genes coding for a MazF cut-site also showed reduced expression in the wild-type compared to genes with no co-cistronic MazF cut-site (Fig. 2c). This was mostly apparent for genes sitting immediately adjacent to the mazF cut-site coding gene (Fig. 2b,c). To directly explore co-cistronic effects, we added an mCherry reporter either upstream or downstream of the GFP_0/3_ reporters in a co-cistronic structure (Fig. 2d). Indeed, comparing the expression of the mCherry reporter in wild-type and Δ*mazEF* backgrounds after 14 hours of growth (deep stationary), showed that the presence of MazF cut-site in the *sfGFP* reporter reduced the expression of both upstream and downstream neighbours, but with a stronger effect on the upstream reporter. Both effects were smaller than the direct effect on the *sfGFP* gene expression. Altogether, the transcriptomic data suggest that the observed effect of MazF on gene expression is driven primarily by direct cleavage and in addition by a co-cistronic effect of the cleavage and through additional downstream effects.

#### Loss of MazEF delays growth recovery and extends the lag phase

Given the strong activation of MazF during the stationary and lag phases of growth and its impact on many target genes, we searched for a phenotype of *mazEF* deletion during the bacterial growth cycle. Wild-type, Δ*mazEF*, Δ*mazF* and the *mazEF* complementation strain were grown in LB medium overnight, diluted 1000-fold and then followed for their growth using a plate reader and characterized for the lag phase duration and subsequent growth rate during the exponential phase (Fig. 3a, Supplementary Fig. S2b, methods). Strikingly, we found that the Δ*mazEF* strain showed a ∼30 minutes increase in the duration of the lag phase, compared to the wild-type (p=0.001, Supplementary Fig. S2a). Δ*mazF* lag was similar to that of Δ*mazEF*, while the complementation strain behaved similarly to the wild-type. No significant effect was found on the growth rate of the cells (Supplementary Fig. S2c,d).

**Figure 3:**
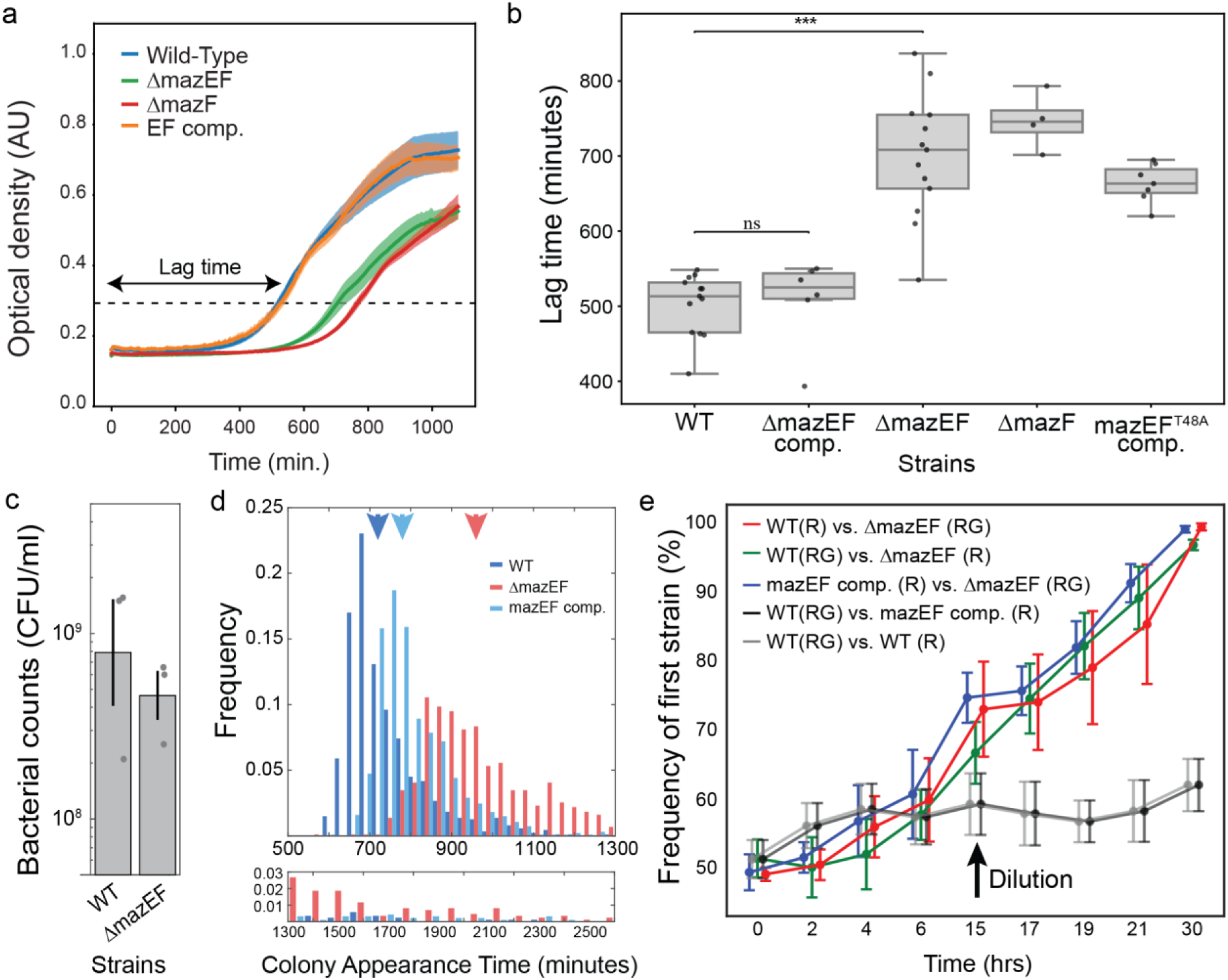
deletion of *mazEF* leads to an extended lag time. (a) An example for growth curves taken on the same day, of different strains. Lag time is set as the duration between dilution into fresh medium to achieving a set OD, as exemplified. (b) Quantification of lag time for different strains. Each dot represents a measurement taken as the mean of three technical repeats. Measurements of the same strain were done on different days. (c) Live cell count after overnight culture of either WT or Δ*mazEF* strains. (d) Distribution of colony appearance times for the different strains, using the ScanLag system. Top: distribution between 500 to 1300 minutes after inoculation. Colored arrows point to the mean of the corresponding distribution. Bottom: continuation of the distribution from 1300 to 2600 minutes. (e) Competition assays for the indicated strains (methods). Shown is the frequency of the first strain in each of the competition types at different times during the growth cycle, as indicated. Error bars represent the standard error for three independent competition experiments. Position of each competition type is shifted from the indicated time for clarity. (R) and (RG) suggest the strains were encoding mCherry, or mCherry+GFP constitutive reporters, to allow frequency measurements during co-culture.

We further characterized the impact of the *mazEF* deletion on growth in minimal medium, and found that Δ*mazEF* extended the lag time by ∼3 hours compared to the wild-type (Fig. 3b,c). In cell cycle terms, the extension of lag time in minimal medium amounts to roughly 3 cell-cycles, compared with ∼1 cell cycle extended lag in LB. As in LB, the phenotype in minimal medium was complemented by ectopic expression of the *mazEF* operon, but not in an operon coding for a non-functional MazF variant (*mazEF*^T48A^, Fig. 3b)(40).

The apparent increased lag after dilution may reflect differences in the initial density of the two strains, rising from different growth/death rates during stationary phase, or from an extended lag time of individual cells of the different strains. Importantly, both CFU counts of overnight cultures and dead/live staining in both LB and minimal medium did not yield significant differences, suggesting that the increased lag cannot be explained by cell death overnight (Fig. 3d, Supplementary Fig. S2e,f). Single cells lag duration and its variability can be masked in liquid cultures by the first cells that resume growth. In order to better characterize the lag of single cells, we used the scan-lag system to monitor the distribution of the time of appearance of colonies of the wild-type and Δ*mazEF* mutant during growth on LB plates (Fig. 3e) (41). While the apparent lag was longer in both strains under these conditions, the results nicely matched the results observed in well-mixed conditions, with the average Δ*mazEF* lag extended by ∼650 minutes compared to that of the wild-type. Both genetic backgrounds showed a unimodal distribution, but the Δ*mazEF* strain had a considerable rightward tail of colonies with extremely long lag (Fig. 3D, bottom). These results suggest that *mazEF* deletion leads to a longer and more variable lag time.

To observe directly the adaptive consequences of the lag difference, we performed competition experiments between a Δ*mazEF* mutant and the wild-type as well as control competition between the wild-type and itself and the wild-type and the complemented Δ*mazEF* strain (Fig. 3e, Supplementary Fig. S3). Competing strains were differentially marked with fluorescent reporters (constitutive mCherry reporter on both strains and constitutive sfGFP on one of the strains) and their relative frequency was monitored by flow-cytometry (methods). We found a substantial increase in the frequency of the wild-type over the Δ*mazEF* mutant during two days of growth-dilution cycles, while no substantial effect was observed in the control competitions. Notably, the greatest increase in wild-type frequency occurred on the second day, after the wild-type exited the lag phase.

#### The MazF regulon is significantly enriched for transcripts governing sporulation and other Spo0A-dependent stress response

Our finding that MazF enable better transition to growth, suggest that it may modulate a stress response occuring during the stationary state. To understand the possible functional role of specific pathways or functions, we searched the transcriptomic data (Supplementary File S1) for enrichment and depletion in specific gene ontology (GO) annotations within groups of genes that were specifically down regulated in the WT compared to all sufficiently expressed genes at the same timepoint (methods). We identified specific enrichment of genes associated with sporulation and related processes (methods, Supplementary Table S1). Notably, the sporulation level at the observed conditions (overnight cultures in LB) is very low, but some sporulation-related genes also control other stress responses(42). Specifically, it was found that in some media, Spo0A activity affects the cellular growth cycle(43).

#### The spo0A stress regulator strongly interact with MazF activity and function during stationary state

The RNAseq experiment pointed to over-representation of sporulation related genes in the MazEF regulon. Sporulation initiation and other related stress responses are guided by the phosphorylated form of the Spo0A master regulator, which is also known to control stationary state physiology(42, 44). We therefore assayed the impact of a *spo0A* deletion on the duration of the lag time in minimal medium and on its regulation by MazEF. Strikingly, we found that the lag time became *mazEF-*independent in a *spo0A* mutant background and slightly shorter than that of the wild-type (Fig. 4A). This epistatic interaction suggests that *mazEF* and Spo0A are incorporated in the same pathway.

**Figure 4:**
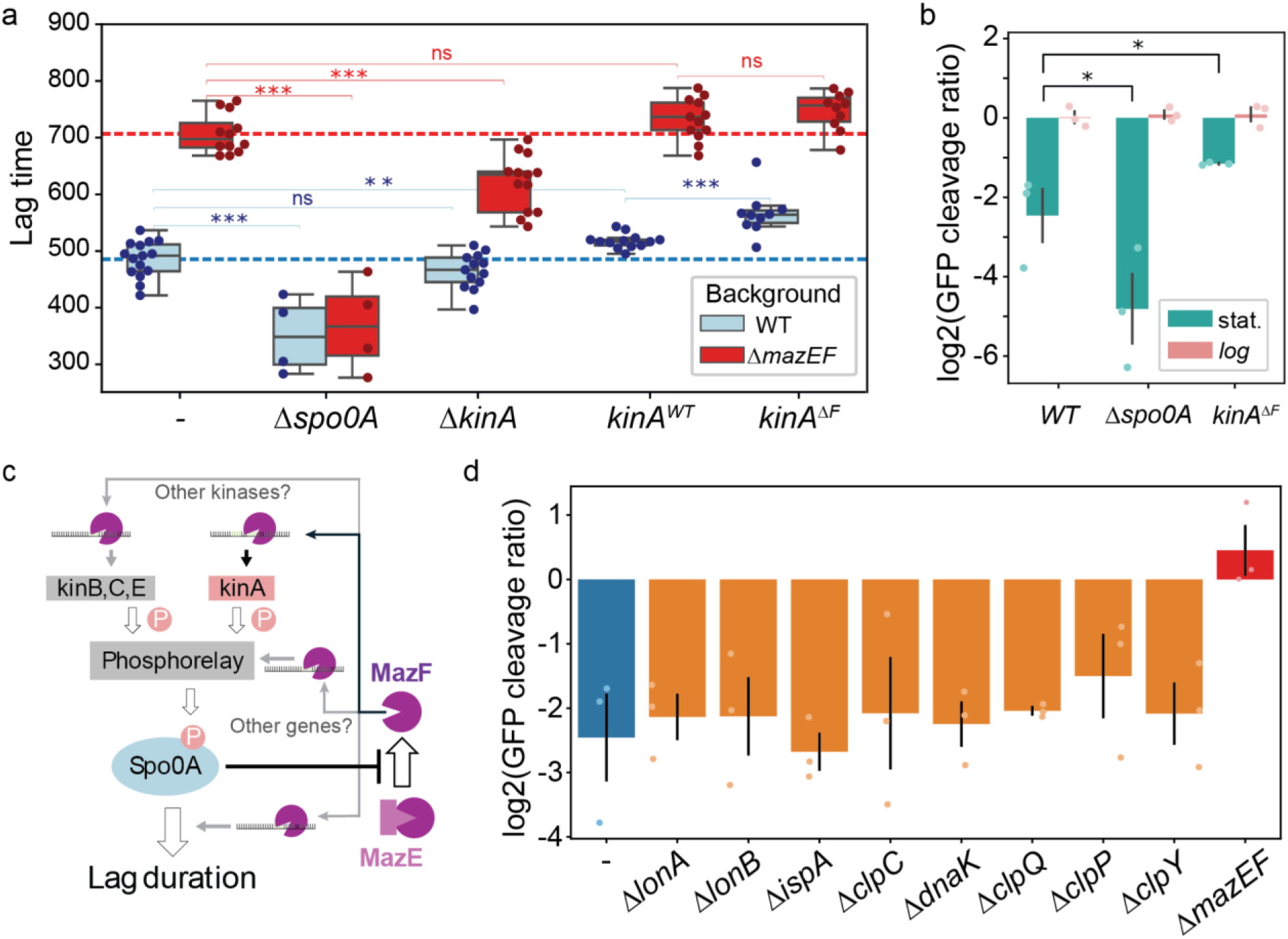
Genetic analysis of Lag time and MazF activity. (a) The lag time in minimal medium is shown for wild-type (blue) and Δ*mazEF* (red) genetic backgrounds with additional genetic perturbations as indicated on the x-axis. kinA^WT^ is a complementation of the kinA deletion mutant, while kinA*^ΔF^* is a complementation of the mutant with kinA recoded to eliminate the two MazF cut-sites. Blue and red dashed lines show the mean lag time of the wild-type and Δ*mazEF* strains for comparison with other strains. (b) GFP cleavage ratio during stationary (green) and logarithmic (red) phases, for different genetic background. (c) Summary of the proposed mechanism identified in (a,b). Black lines mark novel interactions we have identified. (d) GFP cleavage ratio during stationary for different genetic perturbations, as indicated on the x-axis.

One option for the elimination of mazEF-dependence in a *spo0A* deletion mutant is that Spo0A activity, governed by its expression and phosphorylation levels, is necessary for the activation of MazF during the stationary state. Notably, this would not explain the wild-type-like lag phase duration of the mutant, which may be explained by other pleiotropic effects of Spo0A. To examine this option, we assayed MazF activity in a *spo0A* mutant background, using the GFP_0/3_ reporters. Surprisingly, we found the opposite effect - MazF activity during the stationary state became stronger (Fig. 4B,C).

#### Targeted mRNA decay of the KinA kinase mechanistically links MazF activity to lag-phase duration

MazF may affect the lag phase by reducing Spo0A activity or by reducing the expression of related Spo0A regulated genes (Fig. 4C). While *spo0A* itself does not have a MazF cut-site, many of the genes controlling its activity have such sites. Specifically, four out of the five kinases which control Spo0A phosphorylation (KinA,B,C,E but not D) encode a MazF cut-site in their genes, with *kinA* coding for two cut-sites. To study whether MazF cleavage of the kinases can explain the *mazEF* deletion lag time phenotype, we focused on *kinA*, which is known to encode the strongest driver of Spo0A phosphorylation under many conditions(45). Deletion of *kinA* slightly reduced the lag time of the wild-type by 21 minutes (non-significant, p=0.093), but had a stronger effect on the lag time of the *mazEF* mutant, reducing it by 86 minutes (p=5 × 10^−5^) (Fig. 4A). This suggests that KinA contributes to Spo0A phosphorylation at stationary state together with other kinases.

We then complemented the *kinA* mutant by ectopically expressing two different alleles of *kinA* under its native promoter - a wild-type allele (*kinA^WT^*) and a recoded allele (kinA^ΔF^) where the two MazF cut-sites encoded on *kinA* were eliminated (Fig. 4A, methods). We found that the *kinA^WT^* allele over-complemented the *kinA* deletion phenotype in both mazEF^+^ and Δ*mazEF*, leading to a 32 minutes increase in lag time in the wild-type (p=0.002) and 25 minutes increase in the *mazEF* (p=0.08) backgrounds compared to its expression in the native locus. This over-compensation may be due to higher expression at the ectopic location resulting in higher Spo0A activity levels. We then examined the complementation with the MazF-insensitive *kinA^ΔF^* allele. In a Δ*mazEF* background, the *kinA^ΔF^* complementation showed the same lag time as the *kinA^WT^* complementation (p=0.33), suggesting that the sequence modification does not have a *mazEF*-independent effect. Strikingly, complementation with *kinA^ΔF^* in a wild-type (*mazEF*^+^) background significantly extended the lag time by 49 more minutes compared with the *kinA^WT^* complementation (p=3 × 10^−4^). These results suggest that MazF affects Spo0A activity through cleavage of *kinA* mRNA, though this effect explains only around 15% of the total lag-time difference between wild-type and Δ*mazEF*, suggesting that the total effect is multi-factorial. Altogether, these results suggest an intimate linkage between the MazF and Spo0A pathways, which may refine stress-response and the trade-off between stress and return to growth (Fig. 4C).

#### Endogenous MazF activation is independent of major protein quality control proteases

It was previously proposed that the regulation of MazF activation in *E. coli*, may act through degradation of MazE, either by the Clp or the Lon protease complexes (27, 46). Other works suggested that MazF protein levels are dependent on chaperones for proper folding(47). To assess whether these processes also affect MazF activation during stationary state in *B. subtilis*, we examined the impact of deletion of 7 genes coding for components of the main cytoplasmic proteases (LonA, LonB, ClpC, ClpQ, ClpP, ClpY, IspA)(48) and for a major chaperone (DnaK) on the expression of the GFP reporters for MazF activity and found no effect (Fig. 4D). Further work would be required to identify possible additional protein quality control proteins, redundancies masking the effects of the genes examined, or possibly other routes for the activation of MazF (see discussion).

### The 6bp RNA cleavage specificity and regulatory architecture of MazEF are conserved across *Bacillota*

Both *mazE* and *mazF* are core genes of the *Bacillus* genus and the species *Staphylococcus aureus*(35, 49). We therefore wondered what is the scope of conservation of this system in the gram positive (Bacillota) phylum. We extracted all ∼3300 reference sequences of Bacillota species and searched for a gene homologous to MazF (Fig. 5a, Supplementary File S2, methods). We found a homologous MazF protein coded in 80% of these species, with identity levels >60% in large majority of cases. Other species did not show strongly similar proteins. From a phylogenetic standpoint, MazF was found to be completely conserved in many well-studied genera, including *Bacillus*, *Staphylococcus*, *Listeria*, *Clostridium* and *Paenibacillus*, while notably absent from the *Lactobacillus* and Streptococcus genera (Fig. 5a).

**Figure 5:**
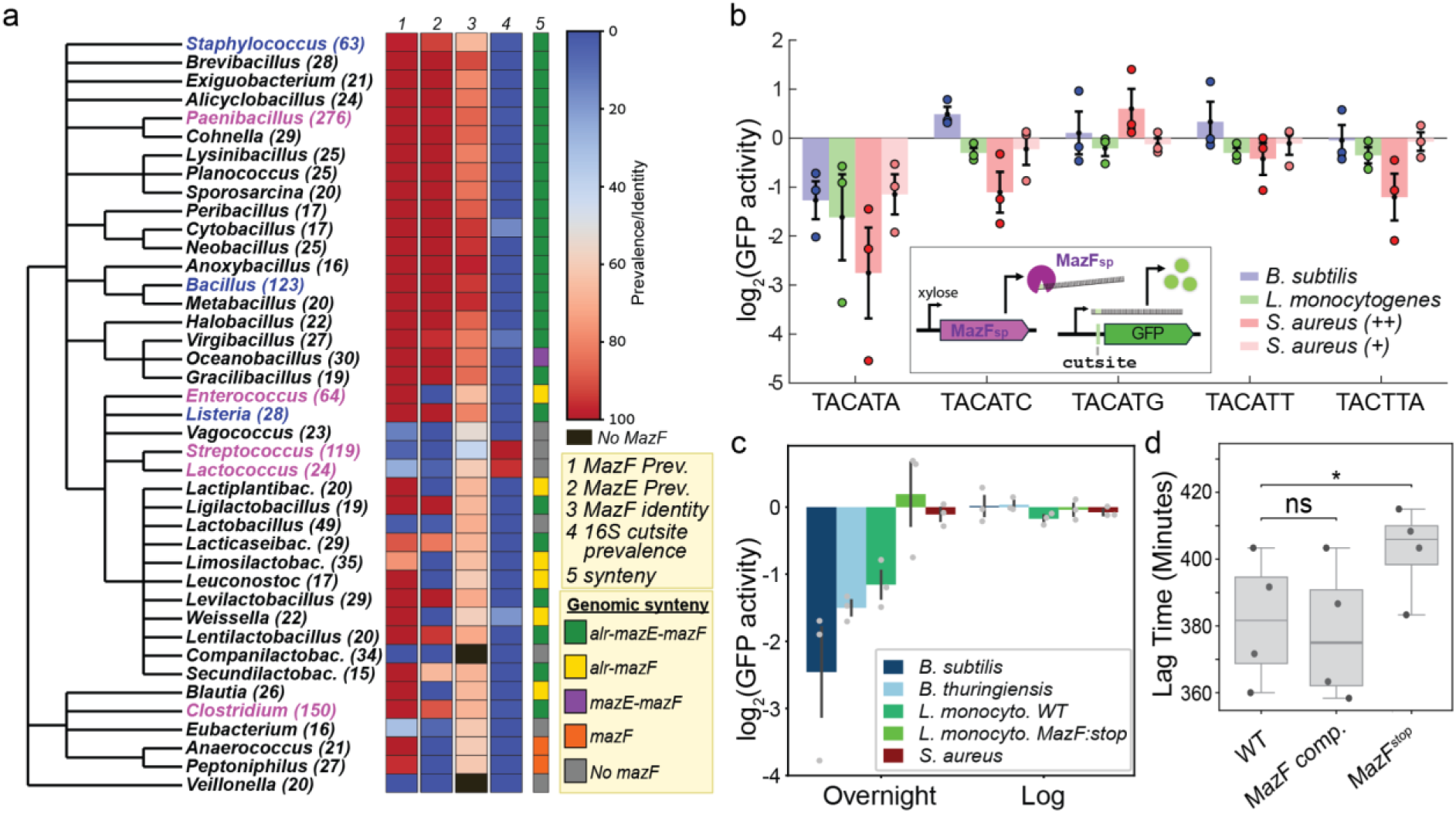
Conservation of MazF and its activity in gram positive bacteria. (a) a taxonomic tree of all genera in the phylum *Bacillota* with more than 15 designated species (methods, Supplementary File S2). For each genus, the number of species it contains is shown in parentheses. Color codes are: 1,2) The percent fraction of species within a genus which code for *mazF* (1) and *mazE* (2) in their genome. 3) The average percent identity of MazF for all species encoding for it within a genus, to MazF of *B. subtilis*. 4) The percent fraction of species within the genus where a TACATA sequence appears in the 16S genes of the reference genome genome. 5) The most common genomic architecture within the genus. Highlighted in blue are the three genera whose MazF activity is measured in panel b,c. Highlighted in magenta are additional genera of interest. (b) Fold change in GFP cleavage ratio, defined here as the expression ratio between a reporter strain induced for MazF with xylose and the same strain when uninduced. Shown are results for five different reporter strains with different putative cut-sites (as indicated on the x-axis) and for three different types of MazF cloned from *B. subtilis* (blue), *L. monocytogenes* (green) and *S. aureus* (red) induced with 0.01% xylose. *S. aureus* was also assayed with induction level of 0.001% (light red) (c) GFP cleavage ratio defined as the expression ratio between GFP_0_ and GFP_3_ introduced into four different species; *B. subtilis*, *B. thuringiensis*, *L. monocytogenes* and *S. aureus*. Shown are results for stationary and logarithmic phases. (d) Lag time of L. *monocytogenes* strains diluted into BHI medium.

In 75% of the genomes with MazF, the upstream gene product to MazF was a putative antitoxin. Notably, the divergence of this protein was much larger where in some cases it was a direct homologue of *B. subtilis* MazE, while in others it did not show sequence homology, but had the same fold or was characterized as a putative antitoxin. In some genomes coding for MazF (eg., in *Enterococci*), no putative antitoxin was found upstream of MazF and it remains open whether there is a non-adjacent negative regulator of MazF in these species as was described in myxococcus (50). The mazEF system (or mazF only in some cases) often appeared in a similar genomic neighbourhood with the *alr* gene upstream of it (Fig. 5a). The synteny of genes downstream of *mazF* was less conserved, though many well-studied genera code for the *rsb* stressosome operon downstream of *mazEF*.

We and others have recently identified the 6bp sequence U^ACAUA as the cut-site sequence for *B. subtilis* MazF (35, 36). Recent works in *S. aureus* recognized two extended cutsite motifs, U^ACAUN and U^ACNUA, while MazF cut-site specificity in other gram-positive systems has not yet been characterized(51). To better probe the conservation of MazF specificity, we constructed sfGFP reporter systems with one of the 5 sequences UACA(U/A/G/C) or UACUUA (matching the second *S. aureus* motif) inserted in the 5’ UTR of the *sfgfp*_0_ gene. We introduced these reporters into a *B. subtilis* Δ*mazEF* mutant and then cloned into each of the reporters a xylose inducible *mazF* gene from either *B. subtilis*, *S. aureus* or *Listeria monocytogenes* (methods). GFP levels were then compared for the 15 strains in the presence and absence of xylose (methods, Fig. 5b). We found that, as expected, MazF_Bs_ cleaved only the UACAUA site and so did MazF_Lm_. MazF_Sa_ showed a stronger reduction in fluorescence for the UACAUA site than that observed for *B. subtilis* or *L. monocytogenes*. It also showed a weak signature of cleavage in the UACUUA and UACAUC sites. By reducing xylose levels, we found that lower induction of MazF_Sa_ led to an observed cleavage effect on the UACAUA site which was similar to that of the other species while cleavage effects on other sites were lost. These results suggest that MazF specificity towards UACAUA sites is well conserved and that other proposed cut-sites are most likely off-targets with reduced affinity.

MazF was proposed to adaptively function in *E. coli* through the cleavage of ribosomal genes (34). In contrast, we find that all ribosomal genes of *B. subtilis* completely lack the UACAUA cut-sites. Extending this to all Bacillota species, we retrieved the 16S rRNA sequence of ∼2200 out of the 3300 reference sequence genomes discussed above. We find that only 3% of the MazF-encoding species had a UACAUA sequence in their 16S gene, compared with 33% of genomes that did not code for MazF (Fig. 5a, Supplementary File S2). While not conclusive, these results suggest that the UACAUA cut-site motif is conserved in other Bacillota species and points towards selection against cleavage of the 16S rRNA by MazF.

#### Population-wide MazF activation during stationary phase is a conserved regulatory strategy in multiple Gram-positive species

We next wondered whether the population wide activation pattern of MazF under stationary state is also conserved in other gram-positive species. To this aim, we introduced a plasmid carrying either GFP_0_ or GFP_3_ reporters into *Bacillus thuringiensis* strain kurstaki hd73, *L. monocytogenes* strain 10403S and *S. aureus* strain JE2. Growing these strains in rich medium, we found that both *B. thuringiensis* and *L. monocytogenes* showed reduced expression of GFP_3_ compared to GFP_0_ during the stationary state, while not showing a difference during logarithmic growth. *S. aureus* did not show a difference in GFP_3/0_ activity in either logarithmic or stationary states (Fig. 5c).

To further explore the function of MazF in *L. monocytogenes* we introduced a stop codon mutation into its *mazF* gene (methods). This led to a loss of MazF activity, as was reflected in the loss of reduction in GFP_3_ levels during stationary state (Fig. 5c). We then compared the lag duration of the wild-type and *mazF* mutant *L. monocytogenes* strains and found that the wild-type lag duration was shorter by ∼1 cell cycle (30 minutes, p=0.012) from that of the mutant. This difference was rescued by complementing *mazEF* from a plasmid (Fig. 5d). Altogether, these results point to a partial conservation of the activation pattern of MazF in gram-positive bacteria and its impact on the lag phase.

## Discussion

In this work, we demonstrate that the MazEF system of *B. subtilis* and other Gram-positive bacteria serves a physiological, non-lethal, regulatory function. While MazF remains inactive during exponential growth, it is uniformly activated upon entry into the stationary phase, where it cleaves a defined regulon of mRNAs coding its highly specific recognition site. By interacting with the Spo0A pathway, this sequence-specific cleavage allows *B. subtilis* to rapidly return to growth upon nutrient replenishment. These structural and functional features are highly conserved across many Gram-positive bacteria. Together, these findings fundamentally challenge the prevailing view of MazEF as a stress-induced toxic switch, supporting a model in which it functions as a post-transcriptional regulator.

### MazF role in the physiology of Bacillus and other gram-positive bacteria

Our work points to a regulatory role of MazF in the normal physiology of *B. subtilis*. Specifically, we have identified its regulatory interaction with the Spo0A pathway (Fig. 4A,B). MazF activity is repressed by Spo0A activation and Spo0A activation is repressed by MazF. We have shown that part of the latter effect is due to MazF cleavage of kinA. The full effect of MazEF is most-likely multifactorial, depending on MazF cleavage of other stress related genes, such as the three additional phosphorelay kinases coding for a MazF cut-site (kinB,C,E).

Spo0A serves as a critical junction in the stress response of *B. subtilis*(44), regulating the sporulation pathway(52), biofilm formation(42) and other stress responses, including the stationary state(43, 44, 53–55). In *B. cereus* group strains, where we have also shown MazF to be activated during stationary phase, Spo0A also contributes to virulence(56, 57). While we have not shown the relation between MazF and Spo0A in those species, we find that despite their considerable divergence(58), phosphorelay kinases of this group are also coding for MazF cut-sites. MazF may therefore also affect Spo0A activity in this group. Whether this is a general regulatory strategy of the Spo0A pathway in spore-forming species remains an open question for now.

While Spo0A-related genes are enriched within the MazF regulon, it also contains many Spo0A-indepenent genes. It is therefore possible that MazF controls other *B. subtilis* stationary stress related processes or other acute responses in a non-lethal manner. Our data also suggest a Spo0A-independent phenotype of MazF in other species. Specifically, *L. monocytogenes* does not code for the Spo0A pathway and MazF regulation of stationary state observed there must therefore rely on other pathways.

### The Logic and Mechanism of Precision RNA Regulation

The native expression of the *B. subtilis mazEF* operon is roughly 100-fold higher than the transcriptional level required to induce an effect on growth when MazF is expressed alone(35). This presumed excess of toxin-antitoxin would have been adequate if MazF was needed to rapidly halt growth. It is less clear what is the adaptive value of this design for transcript regulation. It might be that the excess amount of toxin and antitoxin allows more rapid and quantitative adjustments to MazF activity then would have been possible with lower expression levels or with transcriptional induction of the operon.

Rather than indicating latent toxicity, this excess expression may reflect selection for rapid, reversible, and quantitative regulatory architecture at the RNA level. Regulation at the RNA level may be quicker and less energetically costly under stationary phase conditions.

The endonucleolytic cleavage of transcripts by MazF likely generates unprotected RNA ends, making these fragments optimal substrates for degradation by primary host exonucleases(59, 60). This indirect effect explains the observed reduction in expression in co-cistronic genes neighboring the MazF cut-site enconding genes (Fig. 2). MazF cleavage within a gene most probably leads also to ribosomal stalling on the cleaved site and activation of ribosomal rescue mechanisms, but these have not been studied here.

What leads to the activation of MazF under stationary state? In *E. coli*, it was shown that MazE is degraded by the ClpCP system and it was suggested that rapid degradation can lead to accumulation of free MazF(27, 46). Our analysis of protease mutants did not reveal their involvement in this process (Fig. 4D). However, redundancy between proteases may mask the impact of single gene mutations and further work would be required to eliminate the role of protein degradation in MazF activity. Other mechanisms of MazF activation, such as increased stabilization of MazF or competitive binding to MazE can also lead to MazF activation. The latter process has been recently demonstrated to work during infection of phage phi3T, where host MazF is specifically activated through the competitive binding of SroA and SroB to MazE(38).

### MazF RNA regulation underlying host-phage cross-interaction

The positioning of MazF as a central, highly active transcriptomic regulator provides an evolutionary rationale for its complex interactions with some *Bacillus* bacteriophages, which have been shown to both activate, repress and sense MazF activity(35, 37–39). This activity can provide the phage with information on host physiology, in a similar fashion to the role played by RNAseIII during phage lambda lysis-lysogeny(61). Phage manipulation of MazF activity may also benefit the phage by remodelling host response. Verification of these hypotheses require further study of phage infection at different physiological states.

### The Evolution of Precision in Transcriptome Remodelling by MazF

The 6bp specificity (U^ACAUA) of *B. subtilis* MazF restricts its cleavage to a relatively narrow set of genes (Fig. 2). A 6bp exact-match specificity is exceedingly rare for bacterial toxin ribonucleases or for RNA binding proteins more broadly(62–64), establishing MazF as a highly specific trans-acting RNA factor. This contrasts sharply with the *E. coli* MazF homolog, which recognizes a short ∼3bp motif (^ACA with few additional biases(24, 64)) and targets nearly all mRNAs. In mycobacteria (part of the Actinobacteria phylum), multiple MazF homologs are commonly found and *in-vitro* or over-expression experiments suggested a 4bp motif to some of them(65). However, as we demonstrated in this work (Fig. 5b) and was found by others(51), RNAse overexpression/*in-vitro* experiments can provide lower specificity estimates than what occurs *in-vivo*. It will therefore be important to re-estimate MazF cleavage specificity at in-vivo levels in other phyla to determine their potential as specific mRNA regulators.

Interestingly, *E. coli* MazF has been recently suggested to function non-lethally at low activation levels through the relief of transcription-replication conflicts by cutting 16S rRNA forming R-loops(34). This benefit overcomes a demonstrated cost of non-specific mRNA degradation under conditions of high transcription of 16S genes. Our bioinformatic analysis shows that the ribosomal genes of MazF-coding Gram-positive bacteria specifically lack the UACAUA cut-site (Fig. 5a), pointing to a stark evolutionary divergence in regulatory strategy.

Within gram-positive species, our results demonstrate that MazF is highly prevalent and its high specificity is strongly conserved. This most likely suggest that MazF is utilized as mRNA regulator throughout this phylum, as was verified from our observations of *B. thuringiensis* and *L. monocytogenes*. The changing patterns of response and regulation we observed suggest that the use of sequence-specific mRNA cleavage as a regulatory strategy is conserved in this phylum. This may be deployed in distinct contexts depending on lineage-specific regulatory networks and ecological pressures.

## Acknowledgment

We thank Idan Frumkin, Nitzan Aframian and members of the Eldar lab for comments on the manuscript. We thank Anat Herskovits for help with Listeria strains. This work was supported by grants No. 2288/2021 from the Israel Science Foundation to A.E. Both A.E. and J.P. labs are funded by European Research Council Grant 101118890 (TalkingPhages).

## Author contributions

A.E, R.F. and T.B. conceived this study. R.F, S.O.B, T.B., Y.L. and B.T., S.S, P.G., N.S. and A.Z performed the experiments. T.B performed Data Curation and Analysis R.F, S.O.B, J.P and A.E processed data. R.F. and A.E. wrote the manuscript with inputs from all authors.

## Declaration of generative AI and AI-assisted technologies in the manuscript preparation process

During the preparation of this work the authors used google gemini in order to edit the text. After using this service, the authors reviewed and edited the content as needed and take full responsibility for the content of the published article.

## Data availability statement

Gene expression data and Genomic analysis data are provided as supplementary files to this work

## Methods

### Experimental models

#### B. subtilis

To construct new *B. subtilis* strains, standard transformation protocols were used for genomic integration and plasmid transformation (66). As a genetic background, we have used the *B. subtilis* lab strain PY79 with a markerless deletion *xpf*, which prevents induction of the PBSX prophage derived bacteriocin.

#### L. monocytogenes

To generate *mazF* gene knockout mutants, we introduced 2 sequential premature stop codons at positions 8 and 10, which also introduced the ScaI restriction site used later for mutation validation. Upstream and downstream regions of the gene were amplified using Phusion DNA polymerase and cloned using GeneArt™ Gibson assembly mix (Invitrogen) into the pBHE plasmid (pKSV-oriT). The plasmid was then verified by PCR, and the insert was sequenced. The plasmid was then conjugated to *the L. monocytogenes* 10403S strain cured of its phage, DP-L4056 (Lauer et al., 2002), using *E. coli* SM-10 bacteria. *Trans-*conjugants were selected on BHI agar plates supplemented with chloramphenicol and streptomycin and transferred to BHI medium supplemented with chloramphenicol for two days at 41°C to allow plasmid integration into the bacterial chromosome by homologous recombination. The bacteria were passed several times in fresh BHI medium without chloramphenicol at 30°C to promote plasmid loss and the generation of the mutation. The bacteria were plated on BHI plates with or without chloramphenicol, and sensitive colonies were validated for gene mutation by ScaI restriction of the PCR-amplified product.

GFP coding plasmids were conjugated into *B. thuriengensis* and *L. monocytogenes* using *E. coli* SM-10 bacteria and transformed into *S. aureus* using standard transformation protocols.

Strain construction details, plasmid cloning details and primers are all provided in Supplementary File S3.

### Growth media and conditions

As a rich medium for *B. Subtilis* and *B. thuringiensis* strains in this study, we used LB: 1% tryptone (Difco), 0.5% yeast extract (Difco) and 0.5% NaCl. For the experiments in minimal medium for *B. subtilis* strains, we used Spizizen minimal medium (SMM; 2 g l^−1^ of (NH_4_)_2_SO_4_, 14 g l^−1^ of K_2_HPO_4_, 6 g l^−1^ of KH_2_PO_4_, 1 g l^−1^ of trisodium citrate, 0.2 g l^−1^ of MgSO_4_·7H_2_O), supplemented with trace elements (125 mg l^−1^ of MgCl_2_·6H_2_O, 5.5 mg l^−1^ of CaCl_2_, 13.5 mg l^−1^ of FeCl_2_·6H_2_O, 1 mg l^−1^ of MnCl_2_·4H_2_O, 1.7 mg l^−1^ of ZnCl_2_, 0.43 mg l^−1^ of CuCl_2_·4H_2_O, 0.6 mg l^−1^ of CoCl_2_·6H_2_O, 0.6 mg l^−1^ of Na_2_MoO_4_·2H_2_O); 0.5% glucose served as a carbon source. Liquid cultures were grown with shaking at 220 r.p.m and a temperature of 37 °C. When preparing plates, medium was solidified by adding 2% agar. Antibiotics were added (when necessary) at the following concentrations: spectinomycin, 100 µg ml^−1^; chloramphenicol, 5 µg ml^−1^; kanamycin, 10 µg ml^−1^; macrolide, lincosamide and streptogramin, 3 µg ml^−1^; erythromycin + 25 µg ml^−1^ lincomycin. *Listeria* strains were grown in the rich brain heart infusion (BHI) medium (Merck) at 30 °C. *S. aureus* Bacterial cultures were grown in TSB at 37°C.

### MazF sensitive GFP reporters construction and kinA sites deletion

To construct a reporter for *in-vivo* mazF activity we modified the *sfGfp* DNA sequence in four positions using synonymous mutations that will not alter the original amino acid sequence of the sfGFP protein. pAEC2838 (GFP_1_) was constructed by amplifying the whole pAEC1421 unmodified sfGFP plasmid using the SOB957 and SOB958 primer pair that introduced 2 point mutations, T190C and G192T creating a the first TACATA MazF cleavage site. pAEC2844 (GFP_2_) was constructed by amplifying the whole plasmid pAEC2838 using the SOB959 and SOB960 primer pair that introduced a point mutation A312T adding a second TACATA MazF cleavage site. pAEC2851 (GFP_3_) was constructed by amplifying the whole pAEC2844 plasmid using the SOB961 and SOB962 primer pair that introduced 2 point mutations, T453C and C456A creating a third TACATA MazF cleavage site. Mutants were verified by whole plasmid sequencing.

A similar approach was taken to create the kinA^ΔF^ allele, creating two synonymous mutations resulting in the elimination of the MazF cleavage sites. pAEC3564 was constructed by amplifying the whole pAEC3562 (kinA^WT^ allele) using primers SOB1762, SOB1763 that introduced an A417G point mutation, resulting in TACGTA and eliminating the first MazF cleavage site. pAEC3567 (kinA^ΔF^ allele) was constructed by amplifying the whole pAEC3564 using primers SOB1764, SOB1765 that introduced an A909G point mutation, resulting in TACGTA and eliminating the second MazF cleavage site.

### Growth dynamics

To examine growth dynamics, strains were grown overnight in rich or minimal medium at 37 °C or 30 °C with shaking at 220 r.p.m. then diluted by a factor of 1:1000 into fresh rich media or diluted by a factor of 1:100 into fresh minimal media.

For GFP expression levels, strains were grown overnight in LB diluted then by a factor of 1:100 into fresh LB with xylose 0.01% or 0.001% (w/v), when indicated. OD measurements at a wavelength of 600 nm and GFP expression measurements at an excitation wavelength of 485 nm and emission wavelength 535 nm were performed in a 96-well plate using a plate reader (SPARK multimode microplate reader, Tecan).

### Flow cytometry

Flow cytometry was performed to quantify gene expression at the single-cell level, using a Beckman Coulter Cytoflex flow-cytometer equipped with four lasers (405 nm, 488 nm co-linear with 561 nm, 638 nm). The emission filters used were: mCherry, 620/30, GFP – 525/40.

To determine expression, cultures were grown in LB or SMM overnight at 37° with shaking at 220 RPM and then diluted by a factor of 1:100 into fresh LB or SMM media. At each measurement time-point, 1ml of cultures was separated and IPTG was added to a final concentration of 0.1mM and grown for 1 hour in LB, or 2 hours in SMM and in the overnight cultures prior to flow-cytometry measurement.

For *S. aureus* and *L. monocytogenes* cultures were grown in rich medium overnight at 37° for *S. aureus* and 30° for *L. monocytogenes* with shaking at 220 RPM and then diluted by a factor of 1:100 into media. 1ml of cultures was separated and anhydrotetracycline (aTc) was added to concentration 100 ng/mL. aTc was added and grown for 1 hour prior to the measurement in logarithmic growth, or 2 hours for overnight cultures. Expression levels were measured using flow cytometry.

### RNA-seq experiment

Total RNA was extracted from cells using a High Pure RNA Isolation kit (Roche). To this end, strains were grown overnight in LB at 37 °C with shaking at 220 r.p.m., then diluted by a factor of 1:100 into fresh LB media. Upon reaching OD_600_ of 0.3, cells of wild type, Δ*mazEF*. Samples were centrifuged for 5 min at 2,000 relative centrifugal force, and pellets were flash-frozen in liquid nitrogen. Total RNA was extracted from cells using a High Pure RNA Isolation kit (Roche). Before proceeding to library preparation, RNA quantity and quality were validated via Nanodrop, Qubit and 4200 TapeStation. Ribosomal RNA (rRNA) removal was conducted using the NEBNext rRNA Depletion Kit (Bacteria) (NEB, #E7860L). Paired-end RNA-seq libraries were generated using the NEBNext Ultra Directional RNA kit (NEB, #E7420S). Paired-end sequencing was performed on an Illumina NextSeq500 at the Instrumentation and Service Center at The Weizmann Institute of Science.

### Sequencing read mapping and normalization

Quality of raw fastq sequencing was tested by FastQC software(67). FASTQ files for each barcode were mapped to *B. subtilis* PY79 (accession NC_022898.1) using bowtie2 with the following arguments: -D 20 -R 3 -N 0 -L 20 -i S,1,0.50 -p 6 -I 40 -X 300(68). The samtools suite(69) was used for interconversion of BAM and SAM file formats and conducting indexing. Read densities were calculated by identifying the 5′-and 3′-ends of each paired-end fragment and adding one count position for all positions aligning to and between the paired reads using deepTools (70).

### Analysis of RNA-seq data

To determine the average localized MazF activity around cleavage sites, in the different time points, we used the normalized per base reads of coding regions containing a mazF site with a mean coverage in the WT sample higher than 128 reads across a window of 50 basses upstream and 50 bases downstream of the mazF site and across the three replicates for each time point. We followed closely the protocol presented in ref (64). Briefly, for every basepair, cleavage ratio was determined as the 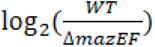 where wt and mutant levels are averaged over the three replicates. Average cleavage ratio as a function of distance from a TACATA cut-site (the C position with negative value for upstream mRNA) was then determined for all cut-sites with expression level with wt coverage level higher than 128 at that condition. Smoothing of the results was performed with scipy signal processing package with savgol filter with a sliding window of 31 and polyorder of 3). As control we used the same approach for 3 different motifs (TACATX where the X is C/G/T).

Gene expression analysis was done using DEBrowser(71). To identify co-cistronic neighbours we used the operons data set from the subtiwiki data set(72). Genes lacking MazF sites that are annotated to be co-transcribed with genes counting MazF sites, were considered as neighboring genes if they were up to 3 open reading frames up/downstream to the MazF-coding gene and were classified as based on their relative position ot the site. In cases that an operon had more than one gene counting a MazF, we used the closest MazF site (in kb) to the gene. Gene ontology (GO) enrichment analyses was done for genes as described in the main text using PANTHER classification system web tool(73).

### CFU count

strains were grown overnight in rich or minimal medium at 37 °C with shaking at 220 r.p.m., strains were diluted 1:10000 and 1:100000 and 100 ul were plated on LB ager plates and grown overnight in 37 °C to allow colonies to form.

#### Live/dead staining

cultures were grown in LB or SMM overnight at 37° with shaking at 220 and were diluted to an OD_600_ = 0.3, Bacterial live/dead viability assays were performed using LIVE/DEAD *Bac*Light Bacterial Viability Kit (Cat. No. L13152).

### ScanLag

strains were grown overnight in rich medium at 37 °C with shaking at 220 r.p.m, strains were diluted 1:100000 and 100 ul were plated on LB ager plates for a CFU of ∼200 per plate. Plates were grown for 48h in 30°C and scanned in 30 minute intervals. Image analysis was then done with ColonyScanalyser python package(74).

#### Competition assays

Constitutive mCherry and sfGFP were used to distinguish between cocultured cells. As the mCherry had an intrinsic cost, we introduced it into both cells and used GFP as the main identifier between strains. LB competition: overnight cultures were diluted by a factor of 100 and grown until they reached OD_600_ = 0.3 and then strains were mixed at a 50:50 ratio, grown and measured for fluorescence at different time-points. Cells were re-diluted after 15 hours. SMM competition: cultures were grown to OD_600_ = 0.1 in SMM and then diluted by a factor of 1,000,000 into fresh SMM and grown until they reached OD_600_ = 0.3. Strains were then mixed at a 50:50 measured for fluorescence. Cocultures were grown overnight and subsequently diluted, regrown and measured for fluorescence at multiple time points.

## Genomic analysis

### Construction of the Bacillota *mazEF* Comparative Database

To characterize the distribution of mazEF systems, we analyzed ∼3,300 reference genome assemblies associated with the phylum Bacillota (NCBI Taxonomy ID 1239). Assemblies were obtained from the NCBI database on July 4, 2024, using the "Reference genomes" filter. To ensure consistent gene prediction across the dataset, open reading frames (ORFs) were predicted ab initio using Prodigal (v2.6.3) in single mode(75). Homologs of the toxin MazF (NdoA), the antitoxin MazE (NdoAI), and the proximal alanine racemase (Alr) were identified using BLASTP-based homology searches(76), using reference sequences from *Bacillus subtilis* 168 as queries. Due to the high sequence divergence typical of antitoxins, we implemented a secondary screening step for operons where MazF was present but MazE was not detected by homology. In these cases, candidate antitoxins were selected based on synteny - specifically targeting the ORF immediately upstream of *mazF*, consistent with the canonical operon architecture. These candidates were analyzed via InterProScan(77) for the presence of the Ribbon-Helix-Helix (RHH) fold, which forms the N-terminal domain of the *Bacillus subtilis* MazE(40). Specifically, candidates were validated as MazE-like if they contained the Arc-like RHH domain (IPR013321), the general RHH domain (IPR010985), or the structurally related SpoVT-AbrB domain superfamily (IPR037914). Results are shown in Supplementary File S2.

### 16S rRNA Extraction and Motif Analysis

Using the genome assemblies compiled for the Bacillota MazEF comparative database, 16S rRNA gene sequences were extracted using custom Python scripts utilizing the Biopython library(78). The extraction pipeline selected optimal representative sequences by prioritizing "16S" annotations with no ambiguous bases and lengths proximal to 1,540 bp. We then quantified the occurrence of the MazF consensus cleavage motif (5’-TACATA-3’) within these sequences. These data were merged with taxonomic lineage information to calculate motif prevalence statistics (mean, median, and maximum counts) across different taxonomic ranks and to identify lineages significantly enriched for the cleavage site. Results are shown in Supplementary File S2.

### Phylogenetic Analysis and Visualization

To visualize these genomic features, a genus-level backbone phylogeny for the Bacillota was derived from the NCBI Taxonomy database. The tree topology was processed using the ETE3 toolkit(79) to prune genera represented by fewer than 15 genomes within the comparative database. The resulting tree was annotated using the Interactive Tree Of Life (iTOL) platform(80). Strain-level data from the comparative database were aggregated by genus to generate iTOL-compatible datasets representing: *mazEF* and *alr* prevalence; mean MazF amino acid identity; dominant operon architectures; and the frequency of the 5’-TACATA-3’ motif in 16S rRNA. To provide statistical context, genus labels were modified to include the total count of analyzed strains.

## Supplementary Figures and Tables

**Table S1:** GO analysis results of over represented pathways in the genes down-regulated in the wild-type compared to *mazEF* deletion. **Bolded** groups are all related to the Spo0A pathway. Analysis was done using the Panther server.

| GO biological process complete | All genes (4260) | Down regulated genes (191) | expected |  | fold Enrich. | P-value | Corrected |
| --- | --- | --- | --- | --- | --- | --- | --- |
| glycogen biosynthetic process (GO:0005978) | 4 | 4 | 0.18 | + | 22.3 | 3.92E-06 | 7.39E-04 |
| glucan biosynthetic process (GO:0009250) | 4 | 4 | 0.18 | + | 22.3 | 3.92E-06 | 6.72E-04 |
| energy reserve metabolic process (GO:0006112) | 5 | 5 | 0.22 | + | 22.3 | 1.72E-07 | 4.64E-05 |
| glycogen metabolic process (GO:0005977) | 5 | 5 | 0.22 | + | 22.3 | 1.72E-07 | 4.06E-05 |
| glucan metabolic process (GO:0044042) | 7 | 5 | 0.31 | + | 15.93 | 3.36E-06 | 7.04E-04 |
| biotin biosynthetic process (GO:0009102) | 9 | 5 | 0.4 | + | 12.39 | 1.87E-05 | 2.94E-03 |
| biotin metabolic process (GO:0006768) | 9 | 5 | 0.4 | + | 12.39 | 1.87E-05 | 2.72E-03 |
| **cellular developmental process (GO:0048869)** | 24 | 7 | 1.08 | + | 6.51 | 5.90E-05 | 7.94E-03 |
| **developmental process (GO:0032502)** | 304 | 39 | 13.63 | + | 2.86 | 7.32E-10 | 1.38E-06 |
| **sporulation resulting in formation of a cellular spore (GO:0030435)** | 278 | 34 | 12.46 | + | 2.73 | 3.79E-08 | 3.57E-05 |
| **anatomical structure formation involved in morphogenesis (GO:0048646)** | 278 | 34 | 12.46 | + | 2.73 | 3.79E-08 | 2.38E-05 |
| **sporulation (GO:0043934)** | 281 | 34 | 12.6 | + | 2.7 | 4.97E-08 | 2.34E-05 |
| **anatomical structure morphogenesis (GO:0009653)** | 283 | 34 | 12.69 | + | 2.68 | 5.95E-08 | 2.24E-05 |
| **anatomical structure development (GO:0048856)** | 283 | 34 | 12.69 | + | 2.68 | 5.95E-08 | 1.87E-05 |

**Figure S1:**
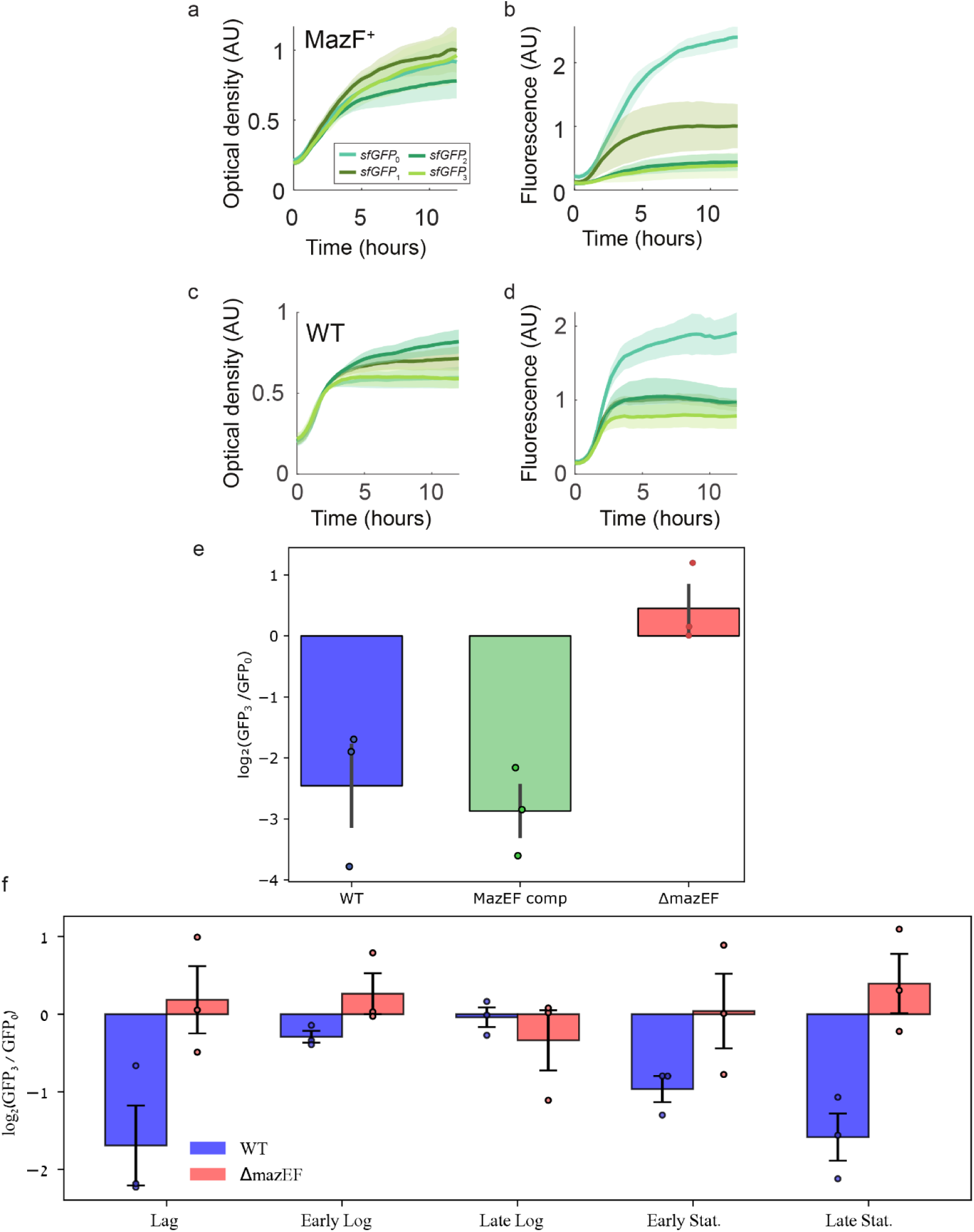
Further analysis of the cleavable GFP constructs. (a-d) Optical density (a,c) and Fluorescence expression (b,d) of strains expressing the GFP_i_ for i=0,1,2,3 in two genetic backgrounds - MazF xylose overexpression (with 0.001% xylose,a,b) and wild-type (c,d). (e) logarithm of GFP cleavage ratio - the ratio in late stationary state between GFP_0_ and GFP_3_ for a wild-type, *mazEF* deletion mutant and a *mazEF* complementation strain (methods). (f) logarithm of GFP cleavage ratio in a wild-type background at different stages of growth in minimal medium (compare to in LB medium in Fig. 1e).

**Figure S2:**
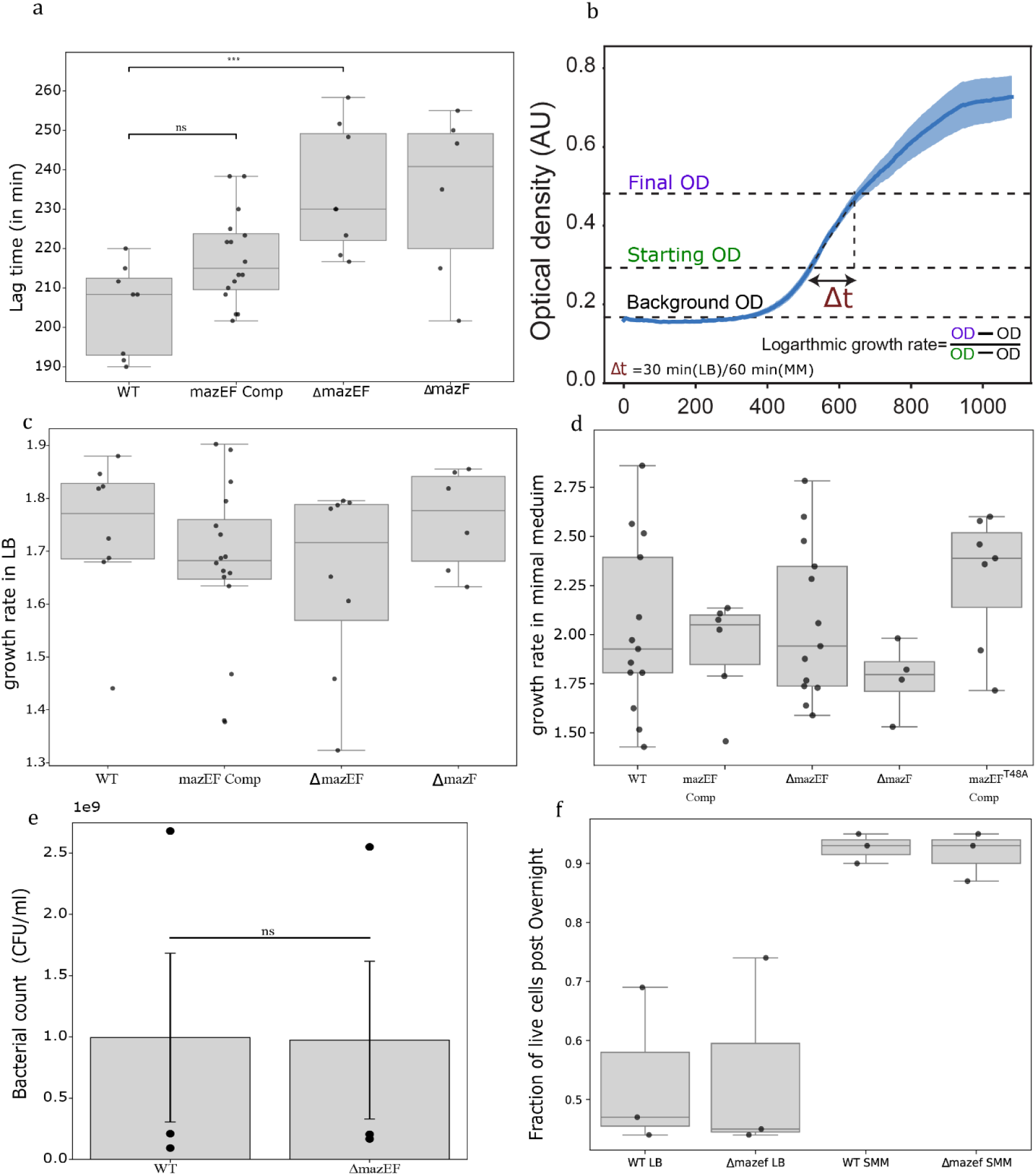
Further information on the growth cycle of wild-type and *mazEF* mutants. (a) lag time duration in LB medium for the indicated strains. (b) Description of the operative measurement of growth rate, exemplified on a wild-type growth curve. Δt is different for measurements in LB and in minimal medium (SMM medium), as indicated. (c,d) growth rate in LB (c) and SMM (d) for different strains. (e) CFU counts for wild-type and mazEF deletion mutant strains grown in LB to deep stationary (15 hours post inoculation). (f) fraction of live cells measured using a live/dead assay (methods) for the two strains in LB and SMM media.

**Figure S3:**
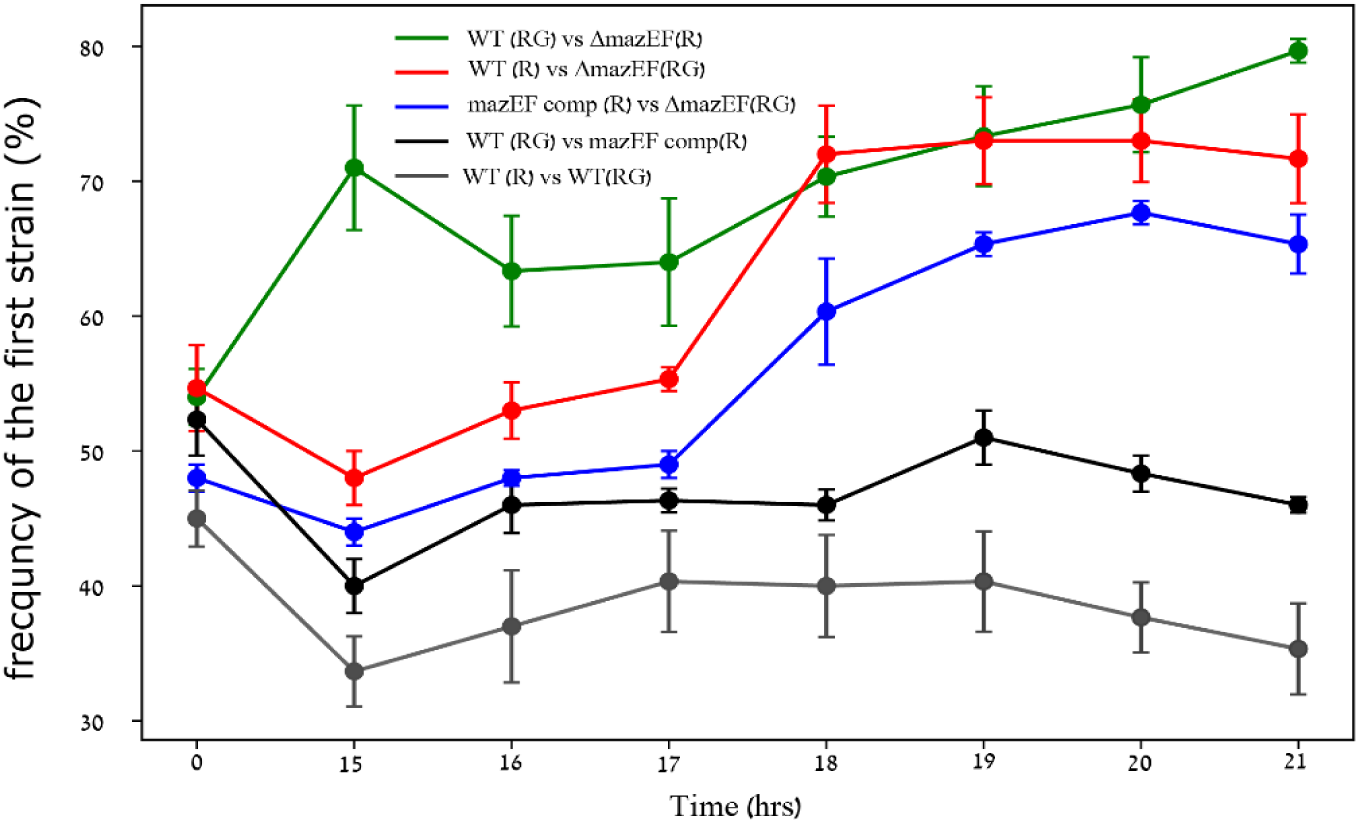
Competition between strains in LB medium. Similar to Fig. 3e, but in LB medium. Note: frequency at timepoints 15,16 hours post dilution are less accurate as RFP expression is low and distinction between live and dead cells is less clear.

## References

1. Qiu, J., Zhai, Y., Wei, M., Zheng, C. and Jiao, X. (2022) Toxin–antitoxin systems: Classification, biological roles, and applications. Microbiol. Res., 264, 127159.

2. Jurënas, D., Fraikin, N., Goormaghtigh, F. and Van Melderen, L. (2022) Biology and evolution of bacterial toxin–antitoxin systems. Nat. Rev. Microbiol., 20, 335–350.

3. Hayes, F. and Van Melderen, L. (2011) Toxins-antitoxins: diversity, evolution and function. Crit. Rev. Biochem. Mol. Biol., 46, 386–408.

4. Harms, A., Brodersen, D.E., Mitarai, N. and Gerdes, K. (2018) Toxins, Targets, and Triggers: An Overview of Toxin-Antitoxin Biology. Mol. Cell, 70, 768–784.

5. Fraikin, N., Goormaghtigh, F. and Van Melderen, L. (2020) Type II Toxin-Antitoxin Systems: Evolution and Revolutions. J. Bacteriol., 202, 10.1128/jb.00763-19.

6. Ramisetty, B.C.M. and Santhosh, R.S. (2016) Horizontal gene transfer of chromosomal Type II toxin–antitoxin systems of Escherichia coli. FEMS Microbiol. Lett., 363, fnv238.

7. Fraikin, N. and Van Melderen, L. (2024) Single-cell evidence for plasmid addiction mediated by toxin–antitoxin systems. Nucleic Acids Res., 52, 1847–1859.

8. Iqbal, N., Guérout, A.-M., Krin, E., Le Roux, F. and Mazel, D. (2015) Comprehensive Functional Analysis of the 18 Vibrio cholerae N16961 Toxin-Antitoxin Systems Substantiates Their Role in Stabilizing the Superintegron. J. Bacteriol., 197, 2150–2159.

9. Wozniak, R.A.F. and Waldor, M.K. (2009) A Toxin–Antitoxin System Promotes the Maintenance of an Integrative Conjugative Element. PLOS Genet., 5, e1000439.

10. Within-host competition selects for plasmid-encoded toxin–antitoxin systems | Proceedings of the Royal Society B: Biological Sciences.

11. Goormaghtigh, F., Fraikin, N., Putrinš, M., Hallaert, T., Hauryliuk, V., Garcia-Pino, A., Sjödin, A., Kasvandik, S., Udekwu, K., Tenson, T., et al. (2018) Reassessing the Role of Type II Toxin-Antitoxin Systems in Formation of Escherichia coli Type II Persister Cells. mBio, 9, 10.1128/mbio.00640-18.

12. Harms, A., Fino, C., Sørensen, M.A., Semsey, S. and Gerdes, K. (2017) Prophages and Growth Dynamics Confound Experimental Results with Antibiotic-Tolerant Persister Cells. mBio, 8, 10.1128/mbio.01964-17.

13. Gerdes, K. and Maisonneuve, E. (2012) Bacterial Persistence and Toxin-Antitoxin Loci. Annu. Rev. Microbiol., 66, 103–123.

14. Page, R. and Peti, W. (2016) Toxin-antitoxin systems in bacterial growth arrest and persistence. Nat. Chem. Biol., 12, 208–214.

15. LeRoux, M. and Laub, M.T. (2022) Toxin-Antitoxin Systems as Phage Defense Elements. Annu. Rev. Microbiol., 76.

16. Kelly, A., Arrowsmith, T.J., Went, S.C. and Blower, T.R. (2023) Toxin–antitoxin systems as mediators of phage defence and the implications for abortive infection. Curr. Opin. Microbiol., 73, 102293.

17. Kwan, B.W., Lord, D.M., Peti, W., Page, R., Benedik, M.J. and Wood, T.K. (2015) The MqsR/MqsA toxin/antitoxin system protects Escherichia coli during bile acid stress. Environ. Microbiol., 17, 3168–3181.

18. Sanchez-Torres, V., Kirigo, J. and Wood, T.K. (2024) Diverse physiological roles of the MqsR/MqsA toxin/antitoxin system. Sustain. Microbiol., 1, qvae006.

19. LeRoux, M., Culviner, P.H., Liu, Y.J., Littlehale, M.L. and Laub, M.T. (2020) Stress Can Induce Transcription of Toxin-Antitoxin Systems without Activating Toxin. Mol. Cell, 79, 280–292.e8.

20. Fraikin, N., Rousseau, C.J., Goeders, N. and Van Melderen, L. (2019) Reassessing the Role of the Type II MqsRA Toxin-Antitoxin System in Stress Response and Biofilm Formation: mqsA Is Transcriptionally Uncoupled from mqsR. mBio, 10, 10.1128/mbio.02678-19.

21. Yamaguchi, Y. and Inouye, M. (2013) Type II Toxin-Antitoxin Loci: The mazEF Family. In Gerdes, K. (ed), Prokaryotic Toxin-Antitoxins. Springer, Berlin, Heidelberg, pp. 107–136.

22. Yamaguchi, Y. and Inouye, M. (2009) Chapter 12 mRNA Interferases, Sequence-Specific Endoribonucleases from the Toxin–Antitoxin Systems. In Progress in Molecular Biology and Translational Science, Molecular Biology of RNA Processing and Decay in Prokaryotes. Academic Press, Vol. 85, pp. 467–500.

23. Zhang, Y., Zhang, J., Hoeflich, K.P., Ikura, M., Qing, G. and Inouye, M. (2003) MazF Cleaves Cellular mRNAs Specifically at ACA to Block Protein Synthesis in Escherichia coli. Mol. Cell, 12, 913–923.

24. Culviner, P.H. and Laub, M.T. (2018) Global Analysis of the E. coli Toxin MazF Reveals Widespread Cleavage of mRNA and the Inhibition of rRNA Maturation and Ribosome Biogenesis. Mol. Cell, 70, 868–880.e10.

25. Mets, T., Kasvandik, S., Saarma, M., Maiväli, Ü., Tenson, T. and Kaldalu, N. (2019) Fragmentation of *Escherichia coli* mRNA by MazF and MqsR. Biochimie, 156, 79–91.

26. Ramisetty, B.C.M., Natarajan, B. and Santhosh, R.S. (2015) mazEF-mediated programmed cell death in bacteria: “What is this?” Crit. Rev. Microbiol., 41, 89–100.

27. Tripathi, A., Dewan, P.C., Ahmed, S. and Varadarajan, R. (2014) MazF-induced Growth Inhibition and Persister Generation in Escherichia coli*. J. Biol. Chem., 289, 4191–4205.

28. Vesper, O., Amitai, S., Belitsky, M., Byrgazov, K., Kaberdina, A.C., Engelberg-Kulka, H. and Moll, I. (2011) Selective Translation of Leaderless mRNAs by Specialized Ribosomes Generated by MazF in Escherichia coli. Cell, 147, 147–157.

29. Alawneh, A.M., Qi, D., Yonesaki, T. and Otsuka, Y. (2016) An ADP-ribosyltransferase Alt of bacteriophage T4 negatively regulates the Escherichia coli MazF toxin of a toxin–antitoxin module. Mol. Microbiol., 99, 188–198.

30. Hazan, R. and Engelberg-Kulka, H. (2004) Escherichia coli mazEF-mediated cell death as a defense mechanism that inhibits the spread of phage P1. Mol. Genet. Genomics, 272, 227–234.

31. Mets, T., Lippus, M., Schryer, D., Liiv, A., Kasari, V., Paier, A., Maiväli, Ü., Remme, J., Tenson, T. and Kaldalu, N. (2017) Toxins MazF and MqsR cleave Escherichia coli rRNA precursors at multiple sites. RNA Biol., 14, 124–135.

32. Guegler, C.K. and Laub, M.T. (2021) Shutoff of host transcription triggers a toxin-antitoxin system to cleave phage RNA and abort infection. Mol. Cell, 81, 2361–2373. e9.

33. Ramisetty, B.C.M., Raj, S. and Ghosh, D. (2016) Escherichia coli MazEF toxin-antitoxin system does not mediate programmed cell death. J. Basic Microbiol., 56, 1398–1402.

34. Fleurier, S. and Matic, I. (2025) MazF endoribonuclease promotes resolution of transcription–replication conflicts at ribosomal RNA genes in Escherichia coli. Nucleic Acids Res., 53, gkaf1034.

35. Guler, P., Bendori, S.O., Borenstein, T., Aframian, N., Kessel, A. and Eldar, A. (2024) Arbitrium communication controls phage lysogeny through non-lethal modulation of a host toxin–antitoxin defence system. Nat. Microbiol., 9, 150–160.

36. Taggart, J.C., Dierksheide, K.J., LeBlanc, H.J., Lalanne, J.-B., Durand, S., Braun, F., Condon, C. and Li, G.-W. (2025) A high-resolution view of RNA endonuclease cleavage in Bacillus subtilis. Nucleic Acids Res., 53, gkaf030.

37. Zamora-Caballero, S., Chmielowska, C., Quiles-Puchalt, N., Brady, A., Gallego del Sol, F., Mancheño-Bonillo, J., Felipe-Ruíz, A., Meijer, W.J.J., Penadés, J.R. and Marina, A. (2024) Antagonistic interactions between phage and host factors control arbitrium lysis–lysogeny decision. Nat. Microbiol., 9, 161–172.

38. Brady, A., Cabello-Yeves, E., Sol, F.G. del, Chmielowska, C., Mancheño-Bonillo, J., Zamora-Caballero, S., Omer, S.B., Torres-Puente, M., Eldar, A., Quiles-Puchalt, N., et al. (2023) Characterization of a unique repression system present in arbitrium phages of the SPbeta family. Cell Host Microbe, 31, 2023–2037.e8.

39. Cui, Y., Su, X., Wang, C., Xu, H., Hu, D., Wang, J., Pei, K., Sun, M. and Zou, T. (2022) Bacterial MazF/MazE toxin-antitoxin suppresses lytic propagation of arbitrium-containing phages. Cell Rep., 41, 111752.

40. Simanshu, D.K., Yamaguchi, Y., Park, J.-H., Inouye, M. and Patel, D.J. (2013) Structural basis of mRNA recognition and cleavage by toxin MazF and its regulation by antitoxin MazE in Bacillus subtilis. Mol. Cell, 52, 447–458.

41. Levin-Reisman, I., Gefen, O., Fridman, O., Ronin, I., Shwa, D., Sheftel, H. and Balaban, N.Q. (2010) Automated imaging with ScanLag reveals previously undetectable bacterial growth phenotypes. Nat. Methods, 7, 737–739.

42. Hamon, M.A. and Lazazzera, B.A. (2001) The sporulation transcription factor Spo0A is required for biofilm development in Bacillus subtilis. Mol. Microbiol., 42, 1199–1209.

43. Zhu, M., Wang, Q., Mu, H., Han, F., Wang, Y. and Dai, X. (2023) A fitness trade-off between growth and survival governed by Spo0A-mediated proteome allocation constraints in Bacillus subtilis. Sci. Adv., 9, eadg9733.

44. Hoch, J.A. (1993) Regulation of the onset of the stationary phase and sporulation in Bacillus subtilis. Adv. Microb. Physiol., 35, 111–133.

45. LeDeaux, J.R., Yu, N. and Grossman, A.D. (1995) Different roles for KinA, KinB, and KinC in the initiation of sporulation in Bacillus subtilis. J. Bacteriol., 177, 861–863.

46. Aizenman, E., Engelberg-Kulka, H. and Glaser, G. (1996) An Escherichia coli chromosomal ‘addiction module’ regulated by guanosine [corrected] 3’, 5’-bispyrophosphate: a model for programmed bacterial cell death. Proc. Natl. Acad. Sci., 93, 6059–6063.

47. Frumkin, I. and Laub, M.T. (2023) Selection of a de novo gene that can promote survival of Escherichia coli by modulating protein homeostasis pathways. Nat. Ecol. Evol., 7, 2067–2079.

48. Harwood, C.R. and Kikuchi, Y. (2022) The ins and outs of Bacillus proteases: activities, functions and commercial significance. FEMS Microbiol. Rev., 46, fuab046.

49. Bukowski, M., Banasik, M., Chlebicka, K., Bednarczyk, K., Bonar, E., Sokołowska, D., Żądło, T., Dubin, G. and Władyka, B. (2025) Analysis of co-occurrence of type II toxin–antitoxin systems and antibiotic resistance determinants in Staphylococcus aureus. mSystems, 10, e00957–24.

50. Nariya, H. and Inouye, M. (2008) MazF, an mRNA Interferase, Mediates Programmed Cell Death during Multicellular Myxococcus Development. Cell, 132, 55–66.

51. Sierra, R., Prados, J., Panasenko, O.O., Andrey, D.O., Fleuchot, B., Redder, P., Kelley, W.L., Viollier, P.H. and Renzoni, A. (2020) Insights into the global effect on Staphylococcus aureus growth arrest by induction of the endoribonuclease MazF toxin. Nucleic Acids Res., 48, 8545–8561.

52. Piggot, P.J. and Hilbert, D.W. (2004) Sporulation of Bacillus subtilis. Curr. Opin. Microbiol., 7, 579–86.

53. Lopez, D., Vlamakis, H. and Kolter, R. (2009) Generation of multiple cell types in Bacillus subtilis. FEMS Microbiol. Rev., 33, 152–63.

54. Nandy, S.K., Bapat, P.M. and Venkatesh, K.V. (2007) Sporulating bacteria prefers predation to cannibalism in mixed cultures. FEBS Lett., 581, 151–6.

55. Molle, V., Fujita, M., Jensen, S.T., Eichenberger, P., González-Pastor, J.E., Liu, J.S. and Losick, R. (2003) The Spo0A regulon of Bacillus subtilis. Mol. Microbiol., 50, 1683–1701.

56. Lücking, G., Dommel, M.K., Scherer, S., Fouet, A. and Ehling-Schulz, M. (2009) Cereulide synthesis in emetic Bacillus cereus is controlled by the transition state regulator AbrB, but not by the virulence regulator PlcR. Microbiology, 155, 922–931.

57. Pettit, L.J., Browne, H.P., Yu, L., Smits, W.K., Fagan, R.P., Barquist, L., Martin, M.J., Goulding, D., Duncan, S.H., Flint, H.J., et al. (2014) Functional genomics reveals that Clostridium difficileSpo0A coordinates sporulation, virulence and metabolism. BMC Genomics, 15, 160.

58. Brunsing, R.L., La Clair, C., Tang, S., Chiang, C., Hancock, L.E., Perego, M. and Hoch, J.A. (2005) Characterization of Sporulation Histidine Kinases of Bacillus anthracis. J. Bacteriol., 187, 6972–6981.

59. Condon, C. (2003) RNA processing and degradation in Bacillus subtilis. Microbiol. Mol. Biol. Rev., 67, 157–174.

60. Mäder, U., Zig, L., Kretschmer, J., Homuth, G. and Putzer, H. (2008) mRNA processing by RNases J1 and J2 affects Bacillus subtilis gene expression on a global scale. Mol. Microbiol., 70, 183–196.

61. Wilson, H.R., Yu, D., Peters, H.K., Zhou, J. and Court, D.L. (2002) The global regulator RNase III modulates translation repression by the transcription elongation factor N. EMBO J., 21, 4154–4161.

62. Lunde, B.M., Moore, C. and Varani, G. (2007) RNA-binding proteins: modular design for efficient function. Nat. Rev. Mol. Cell Biol., 8, 479–490.

63. Bechhofer, D.H. and Deutscher, M.P. (2019) Bacterial ribonucleases and their roles in RNA metabolism. Crit. Rev. Biochem. Mol. Biol., 54, 242–300.

64. Culviner, P.H., Nocedal, I., Fortune, S.M. and Laub, M.T. (2021) Global Analysis of the Specificities and Targets of Endoribonucleases from Escherichia coli Toxin-Antitoxin Systems. mBio, 12, e02012–21.

65. Zhu, L., Phadtare, S., Nariya, H., Ouyang, M., Husson, R.N. and Inouye, M. (2008) The mRNA interferases, MazF-mt3 and MazF-mt7 from Mycobacterium tuberculosis target unique pentad sequences in single-stranded RNA. Mol. Microbiol., 69, 559–569.

66. Harwood, C.R. and Cutting, S.M. (1990) Molecular biological methods for Bacillus Wiley.

67. Andrews, S. FastQC: A Quality Control Tool for High Throughput Sequence Data. Babraham Bioinforma. 2010.

68. Langmead, B. and Salzberg, S.L. (2012) Fast gapped-read alignment with Bowtie 2. Nat. Methods, 9, 357–359.

69. Li, H., Handsaker, B., Wysoker, A., Fennell, T., Ruan, J., Homer, N., Marth, G., Abecasis, G., Durbin, R., and 1000 Genome Project Data Processing Subgroup (2009) The Sequence Alignment/Map format and SAMtools. Bioinformatics, 25, 2078–2079.

70. Ramírez, F., Ryan, D.P., Grüning, B., Bhardwaj, V., Kilpert, F., Richter, A.S., Heyne, S., Dündar, F. and Manke, T. (2016) deepTools2: a next generation web server for deep-sequencing data analysis. Nucleic Acids Res., 44, W160–W165.

71. Kucukural, A., Yukselen, O., Ozata, D.M., Moore, M.J. and Garber, M. (2019) DEBrowser: interactive differential expression analysis and visualization tool for count data. BMC Genomics, 20, 6.

72. Elfmann, C., Dumann, V., van den Berg, T. and Stülke, J. (2025) A new framework for SubtiWiki, the database for the model organism Bacillus subtilis. Nucleic Acids Res., 53, D864–D870.

73. Mi, H., Muruganujan, A., Ebert, D., Huang, X. and Thomas, P.D. (2019) PANTHER version 14: more genomes, a new PANTHER GO-slim and improvements in enrichment analysis tools. Nucleic Acids Res., 47, D419–D426.

74. Levin-Reisman, I., Fridman, O. and Balaban, N.Q. (2014) ScanLag: High-throughput Quantification of Colony Growth and Lag Time. J. Vis. Exp. JoVE, 10.3791/51456.

75. Hyatt, D., Chen, G.-L., LoCascio, P.F., Land, M.L., Larimer, F.W. and Hauser, L.J. (2010) Prodigal: prokaryotic gene recognition and translation initiation site identification. BMC Bioinformatics, 11, 119.

76. Camacho, C., Coulouris, G., Avagyan, V., Ma, N., Papadopoulos, J., Bealer, K. and Madden, T.L. (2009) BLAST+: architecture and applications. BMC Bioinformatics, 10, 421.

77. Jones, P., Binns, D., Chang, H.-Y., Fraser, M., Li, W., McAnulla, C., McWilliam, H., Maslen, J., Mitchell, A. and Nuka, G. (2014) InterProScan 5: genome-scale protein function classification. Bioinformatics, 30, 1236–1240.

78. Cock, P.J.A., Antao, T., Chang, J.T., Chapman, B.A., Cox, C.J., Dalke, A., Friedberg, I., Hamelryck, T., Kauff, F., Wilczynski, B., et al. (2009) Biopython: freely available Python tools for computational molecular biology and bioinformatics. Bioinformatics, 25, 1422–1423.

79. Huerta-Cepas, J., Serra, F. and Bork, P. (2016) ETE 3: Reconstruction, Analysis, and Visualization of Phylogenomic Data. Mol. Biol. Evol., 33, 1635–1638.

80. Letunic, I. and Bork, P. (2024) Interactive Tree of Life (iTOL) v6: recent updates to the phylogenetic tree display and annotation tool. Nucleic Acids Res., 52, W78–W82.

